# Bayesian modeling of skewed X inactivation in genetically diverse mice identifies a novel *Xce* allele associated with copy number changes

**DOI:** 10.1101/2020.11.13.380535

**Authors:** Kathie Y Sun, Daniel Oreper, Sarah A Schoenrock, Rachel McMullan, Paola Giusti-Rodríguez, Vasyl Zhabotynsky, Darla R Miller, Lisa M Tarantino, Fernando Pardo-Manuel de Villena, William Valdar

## Abstract

Female mammals are functional mosaics of their parental X-linked gene expression due to X chromosome inactivation (XCI). This process inactivates one copy of the X chromosome in each cell during embryogenesis and that state is maintained clonally through mitosis. In mice, the choice of which parental X chromosome remains active is determined by the X chromosome controlling element (*Xce*), which has been mapped to a 176 kb candidate interval. A series of functional *Xce* alleles has been characterized or inferred for classical inbred strains based on biased, or skewed, inactivation of the parental X chromosomes in crosses between strains. To further explore the function-structure basis and location of the *Xce*, we measured allele-specific expression of X-linked genes in a large population of F1 females generated from Collaborative Cross strains. Using published sequence data and applying a Bayesian “Pólya urn” model of XCI skew, we report two major findings. First, inter-individual variability in XCI suggests mouse epiblasts contain on average 20-30 cells contributing to brain. Second, NOD/ShiLtJ has a novel and unique functional allele, *Xce*^*f*^, that is the weakest in the *Xce* allelic series. Despite phylogenetic analysis confirming that NOD/ShiLtJ carries a haplotype almost identical to the well-characterized C57BL/6J (*Xce*^*b*^), we observed unexpected patterns of XCI skewing in females carrying the NOD/ShiLtJ haplotype within the *Xce*. Copy number variation is common at the *Xce* locus and we conclude that the observed allelic series is a product of independent and recurring duplications shared between weak *Xce* alleles.

## Introduction

Although X chromosome inactivation (XCI) was first described in the early 1960s (Lyon 1961; Beutler *et al*. 1962), the genetic influences and molecular mechanisms underlying this phenomenon are still incompletely understood. Embryonic stem cells of female placental mammals undergo random XCI, a process that transcriptionally inactivates one of the two X chromosomes early in development (Avner and Heard 2001; Disteche and Berletch 2015). Subsequent daughter cells carry on the initial decision, forming clusters of cells in which either the maternal or paternal X is actively transcribed. Consequently, female mammals are unique mosaics of parental X chromosome activity. XCI ensures that expression of genes on the X chromosome is functionally equalized with those of males as a form of dosage compensation.

At the epiblast stage, each embryonic cell randomly and independently inactivates one of the parental X chromosomes and locks in its cellular fate (Nesterova *et al*. 2001; Okamoto *et al*. 2004). This random selection occurs at around embryonic day E5.5 (Takagi *et al*. 1982; Rastan 1982), prior to differentiation into the three major embryonic germ layers and when there are 120-250 cells comprising the epiblast (Snow 1977). The inactivated X chromosome (Xi) undergoes major reorganization and becomes condensed and heterochromatic, stabilizing gene repression in subsequent somatic cells (Wutz 2011; Nora *et al*. 2012). Regulation of XCI is carried out in part by *Xist*, a *cis*-acting long noncoding RNA (lncRNA) that is transcribed only from the inactivated X (Xi) (Brown *et al*. 1991). The major X inactivation center (*Xic*) extends across a 450 kb multi-function region containing many elements responsible for the complex molecular cascade orchestrating XCI, including *Xist* and other *cis* elements such as *Tsix* and *Xite* (Lee *et al*. 1996; Cattanach *et al*. 1970; Ogawa and Lee 2003).

The role played by *Xist* is necessary but not sufficient to fully explain XCI, leading researchers to explore the larger landscape of *cis* and *trans* regulators, chromatin modifiers, and protein complexes that may comprise the *Xist* interactome (Dossin *et al*. 2020; Penny *et al*. 1996; Brockdorff *et al*. 1991; Giorgetti *et al*. 2016; Minajigi *et al*. 2015). Control of XCI is inherently genetic and thus heterogeneity in the genetic architecture of these elements may affect the expression of *Xist* and its antisense counterpart, *Tsix*, leading to disruption of the machinery controlling the counting, choice, and silencing of the inherited X chromosomes. *Xite* is one such example of a region harboring both allelic heterogeneity and intergenic transcription start sites resulting in differential regulation of *Tsix* expression (Ogawa and Lee 2003). In turn, *Tsix* is monoallelically expressed from the active X (Xa) and blocks *Xist* accumulation, ensuring the future Xa (Lee *et al*. 1999a,b).

XCI is ostensibly random, so the *a priori* distribution of maternal and paternal Xa is expected to be 50:50. Nevertheless, non-random biases between mouse lines have been observed for decades (Cattanach and Isaacson 1967; Cattanach 1970; Cattanach *et al*. 1970), leading researchers to postulate that beyond the control of inactivation, preferential skewing for one parental set of X chromosomes over the other may also be under genetic control. Skewing can take two forms. Primary skewing occurs when the parental chromosomes are inactivated in unequal proportion from the outset (Percec *et al*. 2002). Secondary skewing arises as a form of selection: paternal and maternal chromosomes are initially inactivated at random but the embryonic cells carrying them undergo unbalanced rates of replication or death (Minks *et al*. 2008; Takagi 1980). As such, secondary skewing can be advantageous in the event of a beneficial or deleterious mutation being carried on the chromosome inherited from one parent. The hallmarks of secondary skewing also differ in that it can be tissue-specific and occur at any point during development.

Primary skewing in mice has been associated with an allelic series on the X chromosome named the X chromosome controlling element (*Xce*). Five known functional *Xce* alleles have been described from weakest to strongest, *i*.*e. Xce*^*a*^ < *Xce*^*e*^ < *Xce*^*b*^ < *Xce*^*c*^ < *Xce*^*d*^ (Cattanach and Williams 1972; Cattanach and Papworth 1981). Under this paradigm, X chromosomes with *Xce*^*a*^ are the least likely to remain active, and when found in female heterozygotes alongside the *Xce*^*c*^ allele, skewing as extreme as 20:80 is expected (Figure 1). These allelic designations are well-recognized and have been consistently observed in inbred mouse strains exhibiting replicable skews in X inactivation ratio.

**Figure 1.**
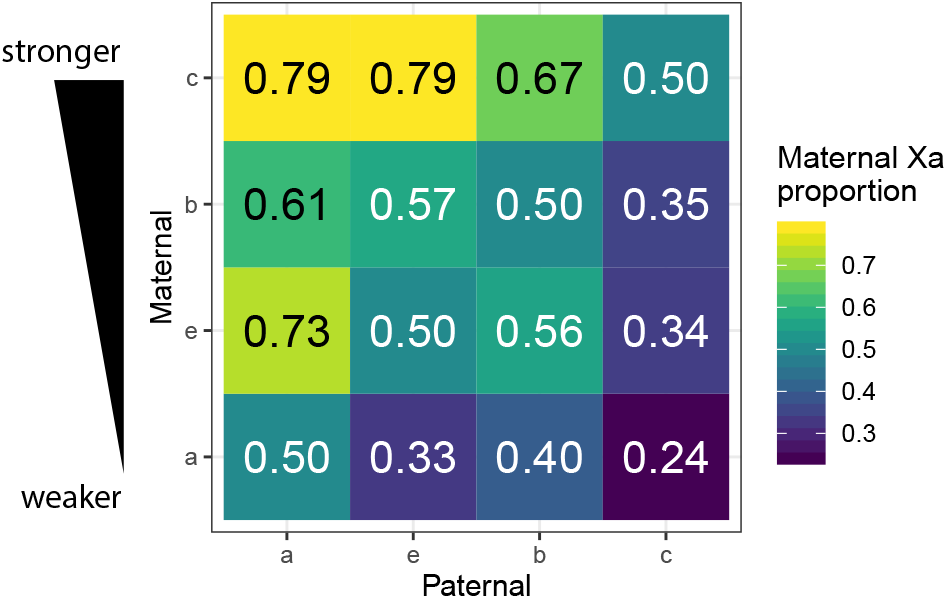
Estimate of active maternal X (Xa) proportion given parental *Xce* alleles from previously published work (Calaway *et al*. 2013; Sheedy 2012; Thorvaldsen *et al*. 2012; Wang *et al*. 2010; Chadwick *et al*. 2006; Plenge *et al*. 2000)

Localizing the *Xce* has required a different set of tools than for a typical quantitative trait locus (QTL). Unbiased localization strategies such as QTL mapping or variant association are optimal for localizing simple, additive QTL. The *Xce* QTL, by contrast, is both multiallelic and completely overdominant in that effects are observable only in the heterozygous state (Cattanach 1970). Those properties, along with the inherent noisiness of the XCI skew trait, rule out the use of such unbiased strategies due to a severe lack of power. Studies localizing *Xce* have therefore tended to be biased towards first principles and focused on the X chromosome: it is logical, mechanistically, that *Xce* must act in *cis* and that there is some distinguishing element the X chromosome so differential preference for one functional allele over the other can manifest.

A natural starting place to search for the *Xce* was within the *Xic*. Control of XCI was initially mapped to a genomic region which overlapped the *Xic*, and *Xist* was an early candidate for the *Xce*. Using translocated coat color genes, Cattanach and collaborators placed the control region for X-chromosome skew between two markers for Tabby (Ta) and Mottled (Mo) coat colors (Figure 2) (Cattanach *et al*. 1969; Cattanach 1970; Cattanach *et al*. 1970). Upon discovery of *Tsix* and *Xite*, allelic heterogeneity across the *Xic* was suggested as a candidate for *Xce* and as an explanation for the phenotypic breadth of skewing observed in mice (Ogawa and Lee 2003). More recent work in the last two decades, however, demonstrated that the *Xce* does not overlap the *Xic*, suggesting that another separate region also participates in XCI. Further refinements over the the decades (Cattanach and Papworth 1981; Simmler *et al*. 1993; Chadwick *et al*. 2006; Calaway *et al*. 2013) have narrowed this down to a 176 kb minimum interval about 500 kb proximal to *Xic*, rich with multiple structural variants including duplications and inversions. Although two studies have suggested XCI skew may be additionally be affected by other regions on the X chromosome (Thorvaldsen *et al*. 2012) and elsewhere (Chadwick and Willard 2005), those effects have not been replicated and no other such regions have successfully mapped. Our working assumption is therefore a single critical region, located in the historically defined interval supported by multiple studies.

**Figure 2.**
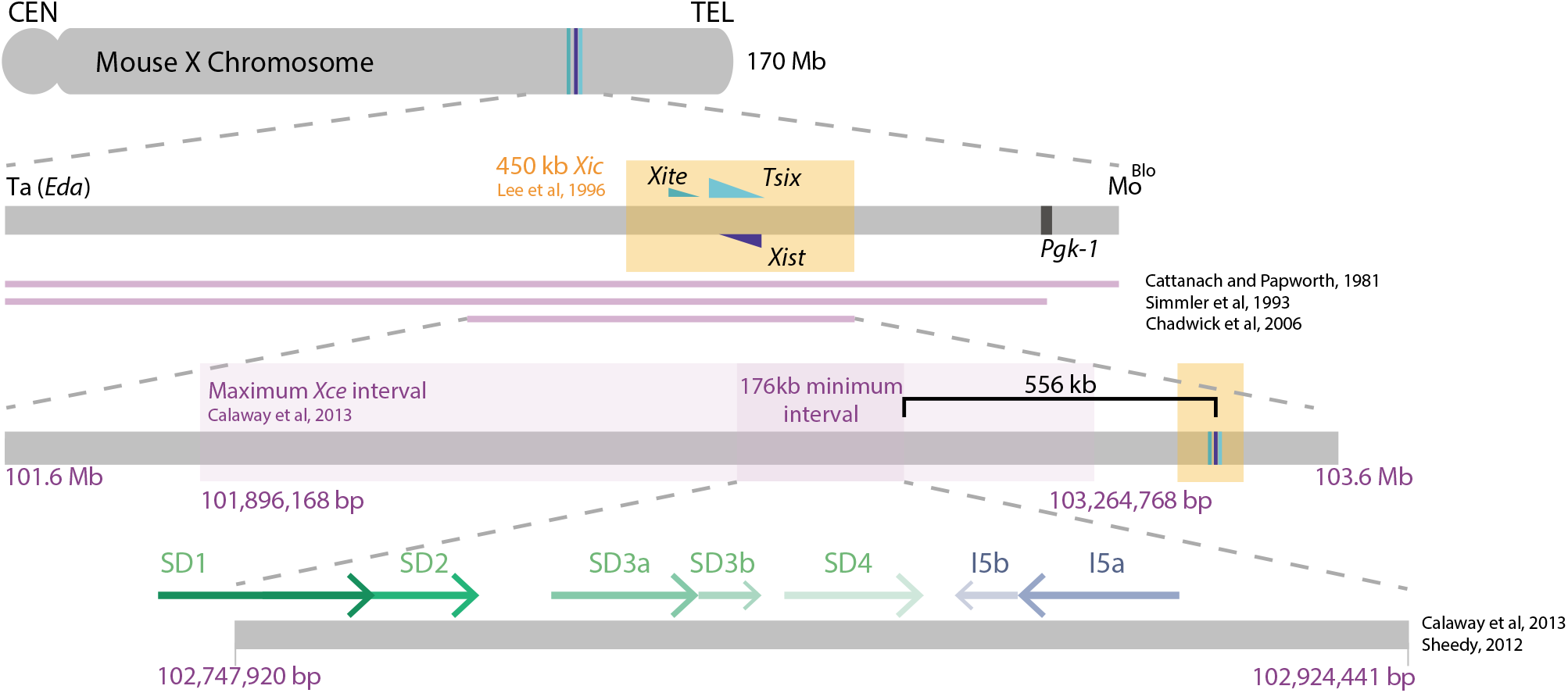
Initial physical mapping and chronological refinement of *Xce* location on mouse X chromosome. Zoomed in region (bottom) depicts the segmental duplications (SD) and inversions (I) examined in this study.

The narrowest proposed *Xce* region to date was reported by our group in Calaway *et al*. (2013) using F1 crosses of classical inbred mouse strains, wild-derived strains, and other *Mus* species. Those results showed that the *Xce* region, localized to an at-minimum 176 kb candidate region consistent with previously described intervals, confers skewed XCI in patterns compatible with the known paradigm (Chadwick *et al*. 2006). The minimum *Xce* interval comprises a series of duplications and inversions and Calaway *et al*. (2013) hypothesized that copy number variations (CNVs) may play a role in XCI skewing (Figure 2). Increased genetic diversity made possible the discovery of another functional allele in the series, *Xce*^*e*^, observed in inbred PWK/PhJ mice (Calaway *et al*. 2013; Crowley *et al*. 2015; Lenarcic *et al*. 2018).

In this study we take advantage of the fairly narrow *Xce* interval defined in Calaway *et al*. (2013) to investigate how sequence and structural variation affects XCI in a genetically diverse mouse population. We further define and characterize the role of *Xce*, and in particular of CNVs, in XCI skewing using 266 female mice from 28 F1 crosses of the Collaborative Cross (CC) multiparental mouse population (Collaborative Cross Consortium 2012; Srivastava *et al*. 2017). The CC are a panel of replicable and genetically diverse inbred mouse strains, each derived from an independent cross of eight inbred strains representing the three major *Mus musculus* subspecies: *domesticus* (A/J, C57BL/6J [B6], 129S1/SvImJ [129S1], NOD/ShiLtJ [NOD], NZO/HlLtJ [NZO], WSB/EiJ [WSB], *castaneus* (CAST/EiJ [CAST]) and *musculus* (PWK/PhJ [PWK]). Each CC strain possesses genome-wide contributions from the founder strains due to mixing that occurred during rounds of breeding, leading to functional genetic variation and phenotypic breadth. Generations of sib-pair mating resulted in inbred haplotype blocks, allowing for replicates of each CC strain.

Most previous studies quantifying XCI have made use of either 1) F1 hybrids of classical inbred mouse strains, or 2) back-crossed mouse populations on an inbred background with specific and deliberate introductions of one other strain to probe the boundaries of *Xce*. In our study, the increased heterozygosity in the genetic background of our CC-derived sample population allows us to tease apart the effects of *Xce* independent from the genetic background. As a result, any observed XCI will be directly attributable to primary skewing due to *Xce* because other loci on the X chromosome will be shuffled among the crosses. Increased genetic heterogeneity in our sample population also allows us to describe further phenotypic heterogeneity in XCI ratios beyond the known *Xce* alleles (Figure 3).

**Figure 3.**
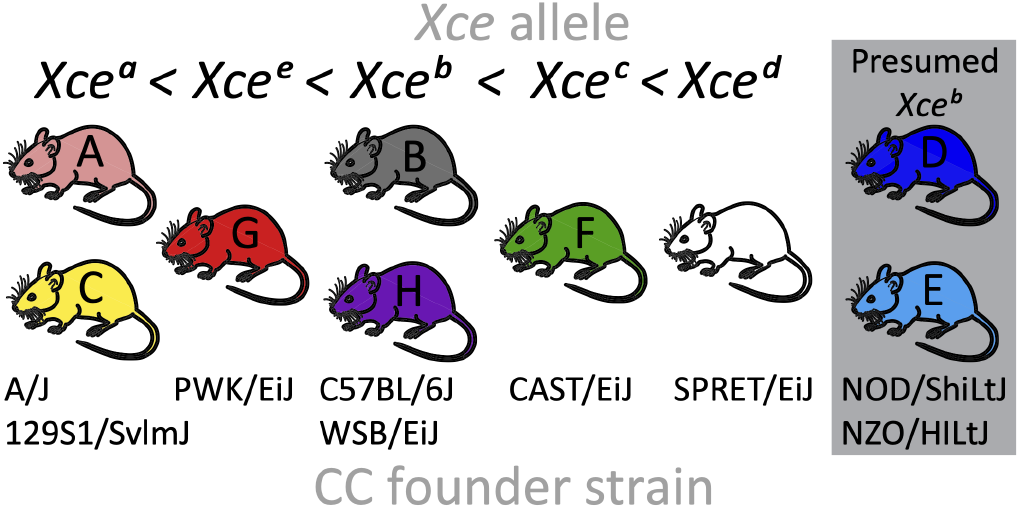
*Mus musculus* strains and their observed or predicted *Xce* alleles. CC founder one-letter code and color corresponds to CC labeling convention. Alleles are ordered in terms of strength. NOD and NZO are presumed *Xce*^*b*^.

Two of the inbred laboratory strains used in generating the CC, NOD and NZO, have not had their *Xce* alleles characterized through crosses; both were predicted to be *Xce*^*b*^ due to haplotype similarity with B6 based on dense genotyping (Calaway *et al*. 2013). Our results interrogate the validity of these predictions based on observed XCI skew in F1 females with sequence derived from NOD or NZO spanning the *Xce*.

Our estimation of XCI skewing is more precise and generalizable compared with much of the XCI literature for two reasons. First, we incorporate X chromosome-wide expression data by quantifying from global RNA-seq. Previous work, by contrast, has generally quantified XCI using allele-specific expression (ASE) measured at a few known genes, which may present biases and inaccurate ratios due to inactivation escape, *cis* regulatory elements, or various confounding variables that are not due to XCI itself. Second, we report precise measures of uncertainty about our estimates using a Bayesian hierarchical statistical model that accounts for multiple sources of information. Chromosome-wide ASE data presents more opportunities for sophisticated statistical modeling to assess XCI, and there are relatively few examples of XCI proportion modeled hierarchically as a beta-distributed random variable (Larson *et al*. 2017; Lenarcic *et al*. 2018). This allows us to largely account for other subtle factors that are known to play a role, such as parent-of-origin effects (POE) in XCI whereby the paternal X (Xp) is predisposed to slightly lower levels of activation regardless of *Xce* allele (Wang *et al*. 2010; Calaway *et al*. 2013; Lenarcic *et al*. 2018). The model also, in accounting for variability in XCI among genetically identical individuals, estimates the effective number of epiblast cells at the point of X inactivation that contribute to the organ on which the RNA-seq is collected.

Another key resource we take advantage of is recently-published high coverage whole genome sequences of the CC strains (Srivastava *et al*. 2017; Shorter *et al*. 2019), which we used to specifically and accurately quantify CNVs across the *Xce*. By quantifying targeted, short reads, we confirm that this region hosts highly recurring sequences which appears to have implications for *Xce* function, and consequently, skewed XCI proportions in mouse crosses. Our characterization of the *Xce* region utilizes the most genetically diverse mouse population to estimate XCI to date and incorporates data from next-generation sequencing to determine ASE, providing a comprehensive quantification of chromosome-wide skewing.

## Materials and Methods

### Notation

Throughout this article, we denote each F1 sample by Strain 1/Strain 2, where counts from Strain 1 comprise the numerator of the XCI proportion, *i*.*e* 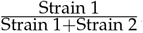. Reciprocal crosses are denoted *a* or *b*, for CC001♀ x CC011♂ and CC011_♂_× CC001 ♀, respectively. These designations were made arbitrarily, but remain consistent throughout the study. Table S1 provides a summary of the CC strains and the F1 crosses.

### Mouse breeding populations and sample collection

The process of generating CC strains has been previously described in detail by Collaborative Cross Consortium (2012). CC mice were purchased from the Systems Genetics Core Facility (SGCF) at the University of North Carolina (UNC). This study includes data from 266 samples derived from a total of 29 CC strains (Figure 4) used to produce 28 F1 recombinant inbred intercross lines (CC-RIX). Data for this study was generated from two CC-RIX sample populations (SP). Heterozygosity present in the RIX lines allows us to both precisely measure ASE by comparing the expression of transcripts with allele A versus transcripts with allele B from mice that inherit the genotype AB.

**Figure 4.**
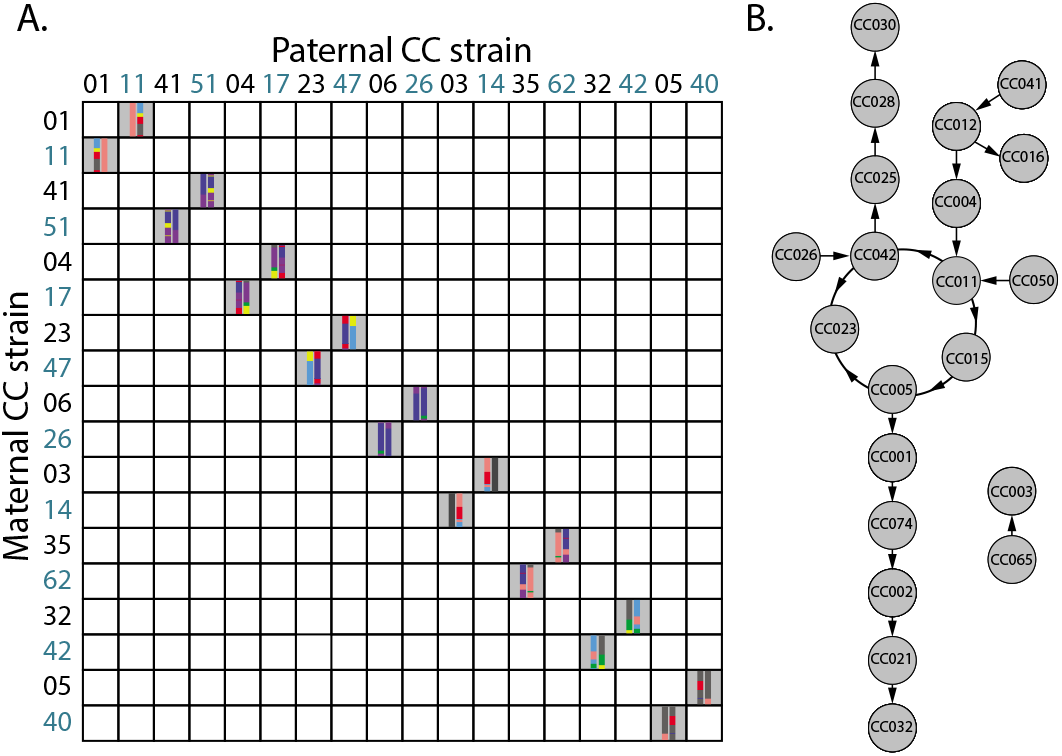
Breeding schemes for two study populations (SP) of CC-RIX mice that contributed data to this study. A) SP1 was developed to study POE, hence the reciprocal RIX design. Schematics of the X chromosome from each founder are shown to illustrate the paired comparisons. B) SP2 provided more diverse pairings of CC strains, without considering reciprocity. CC strains were paired in a quasi-loop design to generate dozens of RIX crosses with maximum diversity, of which this figure only shows the pertinent subset with RNA-seq data. Arrows point from the dam to the sire used for the CC-RIX.

*SP1*: This population was developed to identify strain, POE and perinatal maternal diet effects on gene expression and behavioral phenotypes in adulthood by utilizing F1 crosses of CC-RIX and has been described in detail (Schoenrock *et al*. 2018). Nine genetically distinct reciprocal CC-RIX were bred from 18 non-overlapping CC strains such that samples from CC1♀ x CC2♂ and CC2♀ x CC1♂ are each represented (Figure 4a). Strain-pair selection aimed to maximize several criteria, namely the number of known brain-imprinted loci, as defined from Crowley *et al*. (2015) and Williamson *et al*. (2013) that are heterozygous between haplotypes that are identical by descent with NOD and B6 (Oreper *et al*. 2018).

Females from the 18 CC strains were exposed to one of four experimental diets (vitamin D deficient, protein deficient, methyl donor enriched, or standard control chow; Dyets Inc., Bethlehem, PA) during the perinatal period from 5 weeks prior to mating until their pups were weaned 3 weeks after birth. Whole brain tissue was collected from 188 female CC-RIX mice at 60 days of age (65.1 ± 4.8 days (mean and st dev)). Mice used for gene expression studies were behaviorally naïve. Tissue was collected in 26 batches with a minimum of 2 RIX/diet combinations in a batch. Mice were euthanized and whole brain was immediately extracted. A sagittal cut was made to hemisection the left and right hemisphere and tissue was immediately flash frozen in liquid nitrogen and stored at− 80^°^until pulverization. Right brain hemispheres of all samples were pulverized using a BioPulverizer unit (BioSpec Products, Bartlesville, OK).

*SP2*: In the second population, 21 CC strains, 10 of which overlap with the strains in SP1, produced 19 non-reciprocal RIX. These mice were part of a study to elucidate the genetic basis of antipsychotic-induced adverse drug reactions and has been previously described (Giusti-Rodríguez *et al*. 2020). The larger study comprised 840 mice, representing 62 CC strains and 73 RIX lines. The design of the RIX crosses formed a quasi-loop such that each maternal line was also the paternal line for another cross (see 4b). Only 85 female samples with RNA-seq data were relevant to our analysis so the number of replicates from SP2 is smaller than from SP1 with a median of four samples per CC-RIX (range: 2-7).

Starting at 8-weeks of age, the mice were subjected to a 30 day treatment protocol where half were implanted with slow-release haloperidol (antipsychotic drug) pellets (3.0 mg/kg/day) and the other half received placebo. Treated and untreated mice were matched between sexes, RIX cross, cage, and batch. After 30 days of exposure to drug or vehicle at 12 weeks of age, mice were sacrificed by cervical dislocation without anesthesia to avoid effects on gene expression. Complete description of this experiment is provided in an independent manuscript (FPMV, unpublished).

### RNA-seq preparation

*SP1*: For 188 mice, total RNA was extracted from ∼25 mg of powdered right brain hemisphere tissue using Maxwell 16 Tissue LEV Total RNA Purification Kit (AS1220, Promega, Madison, WI). UNC HTSF core performed RNA concentration and quality check using fluorometry (Qubit 2.0 Fluorometer, Life Technologies Corp., Carlsbad, CA) and a microfluidics platform (Bioanalyzer, Agilent Technologies, Santa Clara, CA). RNA-sequencing was performed in three sequencing batches spread out over the course of the two-year collection of brain tissue once 96 samples from F1 CC-RIX offpsring were obtained. There were a median of 20 samples per CC-RIX (range: 12-32), with 3-4 samples per diet and reciprocal direction.

RNA was prepped with the Illumina TruSeq Stranded mRNA protocol for 100 base pair, stranded, single-end reads at the UNC sequencing core. An initial round of RNA-seq was conducted in December 2014 and June 2015 on HiSeq 2500 machines, and quality control (QC) was conducted on the first few batches of RNA-seq output with fastqc/0.11.8. Reads with low “Per base sequence quality” and “Per sequence quality scores” were prioritized for a second library prep. This first round of RNA-seq was followed up with more sequencing in June 2019 on a HiSeq 4000 machine to boost average read depth for each sample. The final data for each sample were subjected to the same QC criteria and combined, for an average of 24.6 million (M) reads per sample (median 17.9 M, range 10.6-109 M). 7 samples were removed due to missing X chromosomes or low read count.

*SP2*: Detailed methods for RNA-seq sample preparation and processing are described in an independent manuscript (FPMV, unpublished). Briefly, RNA was extracted from striatum using the Total RNA Purification 96-Well Kit (Norgen Biotek, Thorold, ON, Canada) and prepared with the Illumina (San Diego, CA) TruSeq Stranded mRNA Library Preparation Kit v2 with polyA selection using 1 µg total RNA as input. Equal amounts of all barcoded samples were pooled, to account for lane and machine effects. Each of the three pools was sequenced on eight lanes of the Illumina HiSeq 2000 for 100 base pair, stranded, single-end reads.

Quality control filtered out lanes with significant issues in terms of duplication level, fraction of mapped reads (using TopHat2) and, after summarizing reads at a gene level, fraction of mapped reads among the reads that were mapped to an exon. We only considered samples that passed 3 cutoffs: filtering by duplication (at most 40% duplication), percentage of mapped reads (at most 25% reads not mapping) and percentage of mapped reads being mapped to a gene (at most 35% not being mapped to a gene). QC procedures also resulted in corrections or discarded samples due to mismatches in labeling for strain and sex. Principle component analysis identified an outlier that was also removed. Another sample was removed due to a missing X chromosome.

Demographic details about the 266 CC-RIX samples across study populations are compiled in File S2.

### Genotyping in CC-RIX and haplotype reconstruction

To ensure accurate phasing of variants, each sample in SP1 was genotyped on the MiniMUGA platform (Sigmon *et al*. 2020). MiniMUGA is an array-based genetic QC platform with over 11,000 probes designed to perform robust discrimination between most classical and wild-derived laboratory mouse strains. Three X0 females from SP1 that were removed from subsequent analysis were confirmed using the MiniMUGA platform, serving as a useful negative control for our ASE quantification methods. Haplotypes corresponding to each CC founder strain were reconstructed using R/qtl2 v0.20 (Broman *et al*. 2019). Genotype-and allele-probabilities for SP2 were inferred from previous genotyping conducted on CC strains and two to four additional animals per strain known to be their most recent common ancestors using the MegaMUGA platform. MegaMUGA comprises up to 77,800 single nucleotide polymorphism (SNP) markers that were optimized for detecting heterozygous regions and discriminating between haplotypes in homozygous regions, with a special emphasis for markers that are informative in the CC (Morgan *et al*. 2016). Genotyping for MiniMUGA and MegaMUGA was performed at Neogen (Lincoln, NE). Cross-referencing RIX haplotype regions with known CC and CC founder variants for consistency was particularly important at heterozygous loci where the correct parental inheritance would be critical for determining ASE.

We defined the *Xce* in the data based on previously published intervals because all 8 CC founder strains are represented in every sample, instead of each mouse representing one single strain. In iterative stages we defined *Xce*, first, based on the interval described in Chadwick *et al*. (2006) from 101.6-103.6 Mb, and then, refined to the minimum interval described in Calaway *et al*. (2013) roughly from 102.75-102.92 Mb because the narrower interval was still consistent with both our results from the broader interval and previously observed XCI skews between strains. All base pair positions throughout the manuscript are derived from the Genome Reference Consortium Mouse Build 38 (GRCm38).

### Measuring allele-specific expression (ASE) in F1 females

To detect allele- and chromosome-specific expression, we have developed a novel approach using direct k-mer matching to capitalize on known variants in the sequenced CC and founder mice. Key to this method is set of 25-base virtual genotyping probes created from the forward and reverse complement sequences centered at both reference and alternate variants. The reference sequence was provided by the GRCm38 reference mouse genome, based on B6, and alternate alleles were collected from sequence data of the other 7 CC founder strains obtained from the Sanger Institute’s Mouse Genomes Project (Keane *et al*. 2011).

The variant set was filtered to remove unusually high and low probe-sequence counts occurring in any of the sequenced samples. An initial set of approximately 866,000 genome-wide variants were verified across CC and founder strains and became the anchors for matched pairs of k-mers with either the reference or variant allele in the center base. Roughly 590,000 of these k-mers are present in sequences with the highest transcript support level (TSL1), and of those about 414,000 are unique. We filtered k-mers to exclude those that 1) contain multiple variants, and match to 2) duplicated sequences, 3) patterns that are missing from multiple founder strains, 4) loci close to exon start sites, and critically, 5) multiple genomic locations in any CC strain. Taking these criteria into account, between 40-60% of the remaining variants were usable per chromosome. The remaining 7,957 k-mers on the X chromosome comprise a set of paired 25-mers designed to uniquely identify if a sample contains the reference or alternate allele (File S3). We used the tool msbwt v0.3.0 (run on python/2.7.11) to transform our RIX RNA-seq reads into multi-string Burrows-Wheeler Transform (BWT) formatted files to perform efficient, exact k-mer searches to count instances of each k-mer in the RNA-seq reads, thereby quantifying gene expression corresponding to each CC parent in an allele-specific fashion (Holt and McMillan 2014).

### Statistical modeling of X chromosome inactivation

We designed a Bayesian hierarchical model to estimate X inactivation proportion at the level of the gene, individual, and RIX, based on the RNA data above. The model also, as a byproduct of its use of beta distributions and their connection to Pólya urns, estimates the number of brain precursor cells in the epiblast at the point of X inactivation choice, at around E5.5 (Rastan 1982; Lenarcic *et al*. 2018). This section describes first the model for estimating the XCI proportion associated with a given RIX, and then the estimation of the number of brain precursor cells (hereafter, the day 5 brain precursor count) based both on a given RIX and on all RIXs combined. The main components of the model are summarized in Figure 5, with more detail in Figure S1.

**Figure 5.**
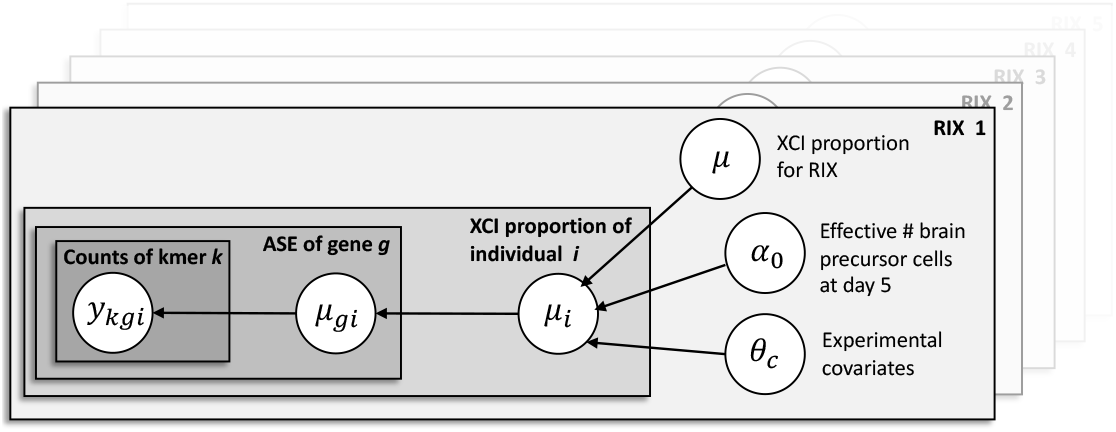
Directed acyclic graph (DAG) showing the main parameters of the hierarchical model for XCI proportion at the gene-, individual mouse-, and RIX level. The *y*_*kgi*_ node is observed; all other nodes are parameters to be estimated. This model is applied to each RIX separately. Estimates for the number of day 5 brain precursor cells (*α*_0_), across RIXs are then combined through a post-processing step.

### Model for RIX-specific XCI proportion

The average XCI proportion inherent to a RIX is reflected by the XCI proportions of mice from that RIX. These mouse-level XCI proportions are in turn reflected by ASE at X chromosome genes. Our model estimates mouse-level XCI proportions for genes by counting k-mers from the allele of one parent vs that of the other and treating these as outputs from a binomial distribution controlled by overall XCI proportions at the gene-, mouse and RIX level.

Consider a given RIX of CC strains *u* and *v*, where strain *u* is expected to have a weaker *Xce* allele or, in the case where both are of the same strength, the maternal strain. For counts associated with *Xist*, which is expressed from the Xi and should therefore have the opposite XCI proportion, the assignment of *u* and *v* were reversed. For mouse *i* = 1, …, *n*, let *N*_*kgi*_ be the total number of counts for k-mer *k* of gene *g* and let *y*_*kgi*_ be the number of these counts specifically from strain *u*. Then, *y*_*kgi*_ is distributed

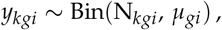

where *µ*_*gi*_ is the expected proportion expressed from strain *u* vs strain *v* for gene *g* in mouse *i*. Different genes *g* = 1, 2, … can have different proportions *µ*_1*i*_, *µ*_2*i*_, …, but we require these to be centered around a common individual-level proportion *µ*_*i*_ as

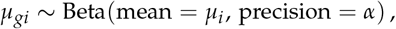

where this corresponds to the conventional parameterization, Beta(*µ*_*i*_*α*, (1− *µ*_*i*_)*α*). The individual-level proportion *µ*_*i*_ is modeled as

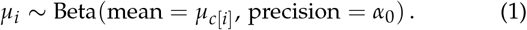

where *c*[*i*] denotes the combination of experimental factors *c* that are relevant to mouse *i, µ*_*c*_ is the XCI proportion predicted for that combination, and *α*_0_ models the day 5 brain precursor count (described later). The proportion *µ*_*c*_ is modeled through a logit link as the outcome of a linear predictor,

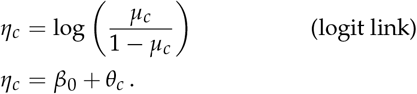

where intercept *β*_0_ models an overall value for the RIX, and *θ*_*c*_ incorporates the effects of experimental covariates.

The set of experimental covariates in *θ*_*c*_ was different for SP1 and SP2. For SP1, these were perinatal diet (diet), POE (recip), and their interaction,

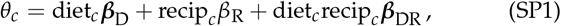

where diet_*c*_ is a categorical predictor indicating the perinatal diet to which mice in condition *c* was exposed, ***β***_D_ is a *n*_diet_ -vector of diet effects constrained to sum to zero, recip_*c*_ indicates reciprocal direction (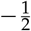 if the dam was *u*, 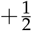 if the dam was *v*), *β*_R_ is the POE, and ***β***_DR_ is a *n*_diet_-vector of treatment-by-POE, also constrained to sum to zero. Across the RIXs in SP1, *n*_diet_ ranged from 2-4, corresponding to a maximum of 4, 6, or 8 conditions per RIX. For RIXs where any condition level *c* contained only one sample, we set *θ*_*c*_ = 0.

For SP2, which did not include reciprocal crosses, we initially considered using

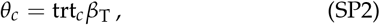

where trt_*c*_ indicates the drug treatment assignment (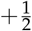 for haloperidol,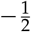 for placebo) of condition level *c*. Treatment assignment was missing for 9 mice, and in these cases we used model-based imputation, 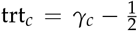 with *γ*_*c*_ ∼ Bin(1, 0.5). The treatment effect, however, was observed to be zero (see File S4), which serves as a negative control for the model given the timing of the drug dose at 8 weeks after birth, well after XCI is established. Because of the zero effect, the lack of a strong biological rationale for its inclusion, and the relative instability of its estimation for some RIX, the final model for SP2 was *θ*_*c*_ = 0, ie, with treatment effect excluded.

Our primary target quantity for each RIX, regardless of its population, was the overall XCI proportion, *µ*, given by the inverse logit of *β*_0_, *i*.*e*.,

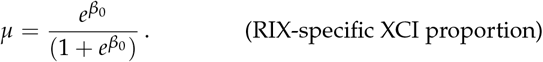

We additionally report XCI proportions for each mouse, *µ*_*i*_ for *i* = 1, …, *n*.

Prior distributions for parameters were specified as follows. For parameters modeling RIX-wide XCI, we set *β*_0_∼ Logistic (0, 1) such that *µ*∼ Unif(0, 1), *i*.*e*., a flat prior on overall XCI proportion. The prior set on *α*_0_∼ Uniform (0, 1000) reflected a reasonable number of cells in the whole embryo at around E5-6 (Snow 1977). Other parameters were modeled with weakly informative priors: *β*_R_, *β*_T_ ∼N(0, 10^4^); ***β***_T_, ***β***_TR_ ∼ N_stz_(**0**, 10^4^× **I**), where N_stz_() is the multivariate normal distribution constrained so that its variates sum to zero [after Crowley *et al*. (2014), Appendix A]; and *α* ∼ Ga(0.01, 0.01).

Posterior distributions for parameters were obtained using Markov chain Monte Carlo (MCMC). MCMC was performed over two separate chains each run with 5 × 10^4^ (SP1) or 10^5^ (SP2) iterations, discarding the initial 10% of the iterations as burn-in and thinning every 5, thus providing 1.8 × 10^4^ or 3.6 × 10^4^ posterior samples in total. Estimates are reported as posterior means (modes and medians are supplied in Tables S1-2) with 95% highest posterior density (HPD) intervals. All models were written and implemented in JAGS 4.3.0 (Plummer 2003) and R version 3.5.2 (R Core Team 2017). Code to run the statistical model is available at https://github.com/kathiesun/XCI_analysis.

### Pólya urn-based estimation of the day 5 brain precursor count

In our model for X inactivation, the individual-specific XCI proportion *µ*_*i*_ is modeled as a beta distribution with precision *α*_0_ (Equation 1). This use of the beta distribution can be directly related to an idealized model of cell proliferation based on a Pólya urn (Lenarcic *et al*. 2018) (Figure S1). The Pólya urn is a hypothetical random process that begins with an urn containing *a* red balls and *b* blue balls. A ball is drawn at random and replaced by two balls of the same color. This is repeated an infinite number of times, after which the proportion of red balls *p*_red_ in the urn will be distributed as

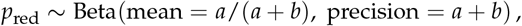

where the precision *a* + *b* is the total number of balls at the point the process began. To the extent that proliferation of embryonic cells in alternate XCI states is analogous to the proliferation of alternate color balls in the Pólya urn, our precision parameter *α*_0_ models the (effective) number of brain-relevant cells at the point of the E5.5 XCI decision.

We estimated 1) an *α*_0_ for each RIX, and 2) a global *α*_0_, based on all RIX data. Posterior distributions of *α*_0_ for each RIX is were obtained using MCMC as described above. These were similar to each other but individually somewhat vague (see **Results**). To obtain a more precise estimate, we assumed the *α*_0_ was the same across RIXs and calculated a posterior given all RIX data as the normalized product of the individual posteriors,

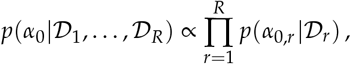

where *p(α*_0, r_ | *𝒟 r*) denotes the posterior for RIX *r* = 1, …, *R* given RIX data 𝒟 _*r*_, and the above relation holding only because the priors on *α*_0_ are identical and uniform such that 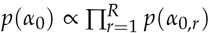. In practice, this involved parametrically approximating each RIX posterior, *p*(*α*_0,*r*_| 𝒟 _*r*_), as gamma distribution with shape *Â*_*r*_ and rate 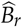 using the `fitdistr()` function from the R package MASS v7.3-51.4 (Venables and Ripley 2002), and then calculating their renormalized product, which is equivalent to a gamma distribution with shape 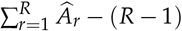 and rate 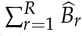.

Point and interval estimates from the aggregate posterior approach above were comparable to those from traditional random-effects meta-analysis on the per-RIX estimates, the latter conducted with the R package `meta` v4.14-0 (Balduzzi *et al*. 2019) using both inverse variance and DerSimonian-Laird estimators (DerSimonian and Laird 1986).

### Whole genome sequences of CC strains

Over the last few years, high-coverage sequences of the CC strains have been made available to the research community. These whole genome sequences (WGS) improved upon the resolution of recombination breakpoints and haplotype assignment in 75 CC strains by sequencing paired-end short reads (150 bp) at 30× coverage for a single male per strain (Srivastava *et al*. 2017; Shorter *et al*. 2019). Deeper sequencing led to improved haplotype reconstruction in samples bred from CC strains, and allowed for identification of unique mutations private to a particular strain. We incorporated additional WGS of the CC founder strains from other previously published sources (Keane *et al*. 2011) and from the GRCm38 mouse reference genome.

The WGS described above for 75 CC strains, along with 24 replicates of B6 mice and one replicate each of the other seven CC founders, have been made publicly available in BWT-formatted DNA-seq reads http://csbio.unc.edu/CEGSseq/index.py. These multi string BWTs were built using the msBWT python tool (Holt and McMillan 2014) from all lanes and paired ends of the Illumina read sets for these genome sequences. Resources making use of the the BWT dataset for effecient k-mer searches have been previously described (Srivastava *et al*. 2017).

### Haplotype analyses based on WGS

The resulting WGS from the CC strains were used to assemble 8 intervals totalling 8,215 bp across the Calaway *et al*. (2013) minimum *Xce* locus in each one of the 8 CC founders. The following CC strains represented the corresponding founder as follows: reference genome for B6; CC055 as representative of the NOD haplotype; CC020 for A/J; CC024 for 129S1; CC051 for WSB; CC032 for CAST; CC003 for PWK; and CC002 for NZO. We first identified intervals between 0.4 – 3 Kb in length, composed of contiguous 45-mers that are present only once in the reference genome. We used the most proximal of these 45-mers as a seed and assembled the sequence in the CC strains using the consensus of the read pileups. All bases used in the consensus were supported by at least two independent reads and, within each strain, lacked any evidence of SNPs or copy number differences. Once assembled, the sequences were aligned using the EMBL-EBI tool, Clustal Omega (Madeira *et al*. 2019), and alignments were optimized by manual inspection to reduce the number of variants. The location, length, and CC strains used for the assembly are shown in Table S3.

### Phylogenetic analysis of CC founder strains

The 8 assembled intervals spanning the *Xce* region were used to estimate the phylogenetic relationship based on X chromosome sequence similarity among the 8 CC founders using BEAST 2.6.3, which performs Bayesian evolutionary analysis by sampling trees (Bouckaert *et al*. 2019). The tree model was based on a coalescent prior for a constant population and was simplified with linked site, clock, and tree parameters among the intervals. We assumed a strict clock and the HKY substitution model (Hasegawa *et al*. 1985). We generated 10^7^ MCMC samples from the posterior of coalescent trees, thinning every 10^3^ samples, over the course of three separate runs with different starting seeds for a total of 3 ×10^4^ recorded posterior samples. We visualized the resulting tree set using DensiTree.v2.2.7, which shows different topographies with varying level of support.

### Quantifying copy number variations

The set of 106 WGS with BWT-formatted data described above was also previously used to develop an occurrence-count matrix of every sequential, non-overlapping 45-mer from the standard mouse reference (GRCm38). We used this count matrix to query 45-mers across CC strains containing different functional alleles in the *Xce* interval defined in Calaway *et al*. (2013). By comparing and quantifying differential k-mer counts between strains, we generated discrete evidence of CNVs in regions along the X chromosome. Samples were classified into eight groups corresponding to the CC founder strains at the *Xce* interval, roughly between 102.65-102.95 Mb when translated to GRCm38 coordinate space. The 24 B6 replicates comprised the baseline “reference” group and the remaining CC-derived samples that were homozygous for B6 across the *Xce* interval were separated into another group to provide a negative control.

Strain-wide copy numbers for each k-mer were first normalized per sample and then averaged across samples in each group. Segmental duplications (SD) and inversions (I) were defined as regions where the mean difference, Δ, between 45-mer counts in the comparison strain versus the inbred B6 mean were different than 0 after k-means clustering of Δ centered at 0, *>* 0, and < 0. K-mers that have an average of one copy in the reference group and zero copies in the comparison group were deemed to contain nucleotide polymorphisms in the non-reference strain. The relevant 45-mers spanning the *Xce* are compiled in File S5, along with the X chromosome positions, the number of copies present in the reference genome, and any SD or I assignments. Alignment boundaries for each SD were determined and visualized using Gepard v1.40 with word size of 45 (Krumsiek *et al*. 2007).

### Data availability

The processed data and code to support the results reported here are available at Figshare (https://figshare.com/s/04b434e49df10ce1343d). These data include: full demographic data for SP1 and SP2; curated lists of 25-mers used to detect reference and variant alleles in RNA-seq data from the X chromosome along with code to generate this list; k-mer counts of the curated 25-mers for both populations; k-mer counts of 45-mers from DNA-seq using CC and CC founder strains; positions of segmental duplications and inversions in the *Xce*. The processed incident count matrices of contiguous 45-mers for the CC strains noted above, and BWT-formatted files of all RNA-seq data are available publicly at http://csbio.unc.edu/CEGSseq/index.py. Genotyping data for the CC MRCAs are available at http://csbio.unc.edu/CCstatus/index.py?run=FounderProbs and genotyping data for SP1 have been deposited in a UNC Data-verse repository (https://dataverse.unc.edu/dataverse/MiniMUGA) under DOI number 10.15139/S3/UYURKF. All R scripts to run the statistical model, and process and generate datasets are available at https://github.com/kathiesun/XCI_analysis.

## Results

### XCI ratio estimated for each mouse and RIX from RNA-seq allele-specific expression

The CC-RIX females comprising this study were genetically heterogeneous mosaics of the 8 CC founder strains with one copy of each chromosome inherited in its entirety from each CC parent. In order to quantify ASE, we relied on efficient multi-string BWT searching of k-mers to identify reference and alternate alleles in the RNA-seq reads. This is akin to a microarray-based quantification strategy where each k-mer represents a probe designed based on prior knowledge, allowing us to precisely target known SNPs to measure ASE.

Counts of k-mers containing reference and alternate alleles of variants were attributed to a particular CC parent according to the haplotype reconstruction derived from genotyping data (Files S6-8). The relative frequency of summed reads across a gene originating from one of the CC parents, *e*.*g*. CC001 in a CC001/CC011 RIX, was modeled analogously to the frequency of heads when flipping a potentially biased coin, as a binomial count that depends on an underlying long-run proportion that may deviate from 0.5. This proportion was estimated for each gene; the proportions across genes were used to estimate an underlying XCI proportion for each mouse; and the XCI proportions across mice were used to estimate a proportion specific to the RIX. These estimations were performed simultaneously using a Bayesian hierarchical model, which also 1) incorporated, and thereby corrected for, potential effects of experimental or breeding-related factors, and 2) connects the variability of mouse-specific XCI proportions about their RIX-wide mean to the subset of epiblast cells at the point of the initial XCI decision contributing to the assayed tissue, in this case the brain.

### XCI is relatively consistent across genes within an individual

Across an individual mouse, gene-level estimates of XCI proportion are stable, suggesting that our quantification methodology is reliable. Figure 6 shows XCI proportion estimates for a mouse each from three CC-RIXs (all 266 samples are shown in File S9). Our Bayesian model estimates posterior distributions for XCI proportions at the gene and RIX level; we report both the means of those distribution and their 95% highest posterior density (HPD) intervals. Gaps in the X chromosome position reflect the patchwork heterozygosity and homozygosity of the CC-RIX samples. In this example, the HPD intervals are fairly narrow around the means, indicating the precision of these estimates, and for two of the mice, the XCI proportion is far from 0.5, indicating strong XCI skew (File S2).

**Figure 6.**
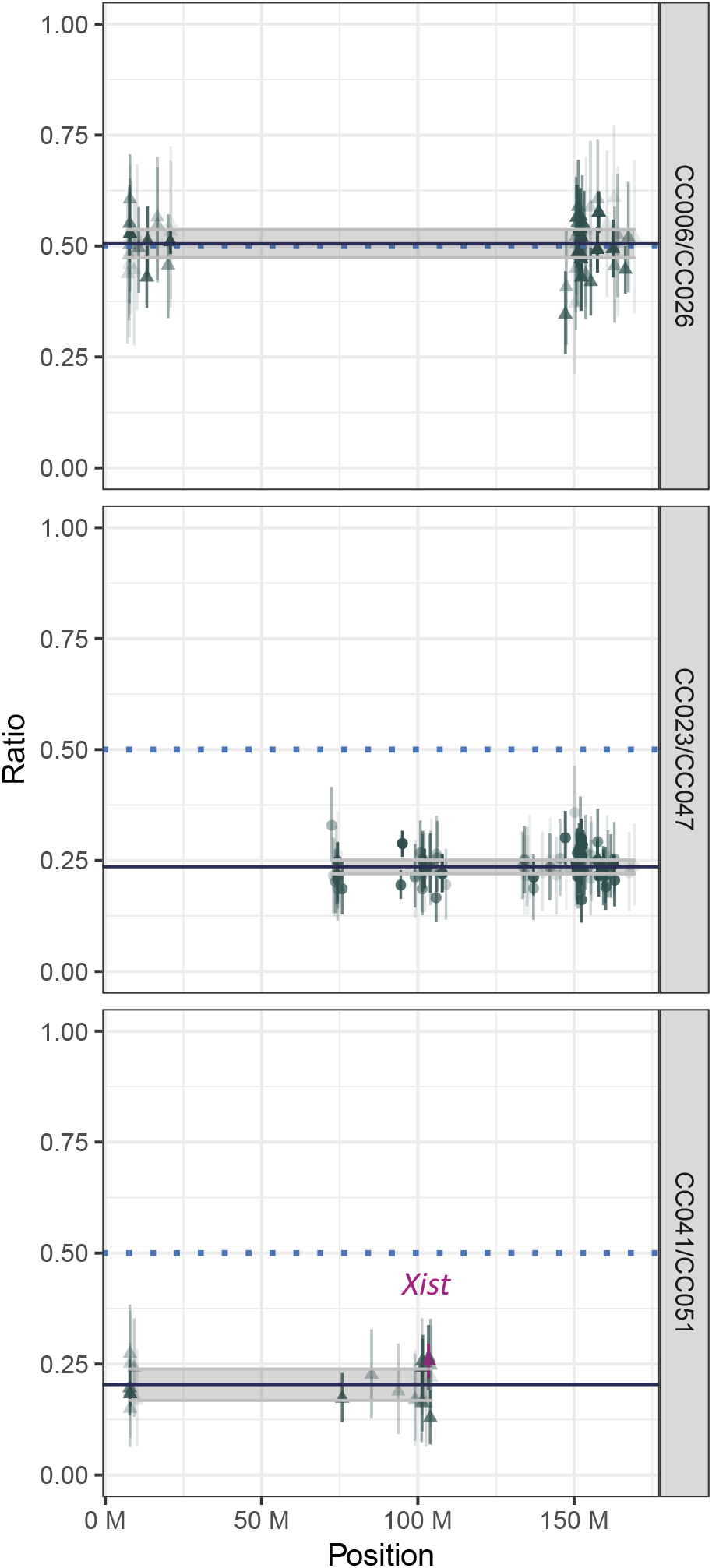
Proportions of parental X chromosome representation at the gene-level (points with 95% HPD bars) for one mouse in each of three separate CC-RIX. Mouse-level estimates summarized as line across the region and shaded 95% HPD interval. Counts from the first strain cross name contribute to the numerator of the proportion. *Xce* allele status: A) CC006 and CC026 are both *Xce*^*f*^; B) CC023 (*Xce*^*f*^) has a weaker allele than CC047 (*Xce*^*b*^ derived from NZO); C) CC041 (*Xce*^*f*^) has a weaker allele than CC051 (*Xce*^*b*^ derived from WSB).

These three example mice demonstrate the consistency of estimates for each sample at genes across the X chromosome, supporting estimates of even fairly extreme XCI skews such as those shown in the Figure 6b-c. At the mouse level, this consistency is representative of samples in the experiment overall.

### Pattern of XCI skew in RIXs with known Xce allele is consistent with previous studies

Our results for XCI skew were largely consistent with previously published research, given our knowledge about the underlying haplotype structure of the CC strains and the known *Xce* subtypes corresponding to major *Mus musculus* strains (Figure 3). Estimates of XCI proportions for each sample and each CC-RIX are compiled in Table S1 and File S2.

Figure 7 shows the XCI proportion at the individual- and RIX-level for every cross in the study, divided as crosses between strains with a) previously phenotyped *Xce* alleles and b) inferred alleles. Crosses with both CC strains sharing the same *Xce* allele had XCI ratios at roughly 50:50. The crosses demonstrate that *Xce*^*a*^ is weaker than any other known allele, as only roughly 30-35% of the cells have active chromosomes bearing *Xce*^*a*^ (Figure 7a). *Xce*^*e*^ and *Xce*^*b*^ are approximately of equal strength, which corroborates the similar pattern seen in Calaway *et al*. (2013).

**Figure 7.**
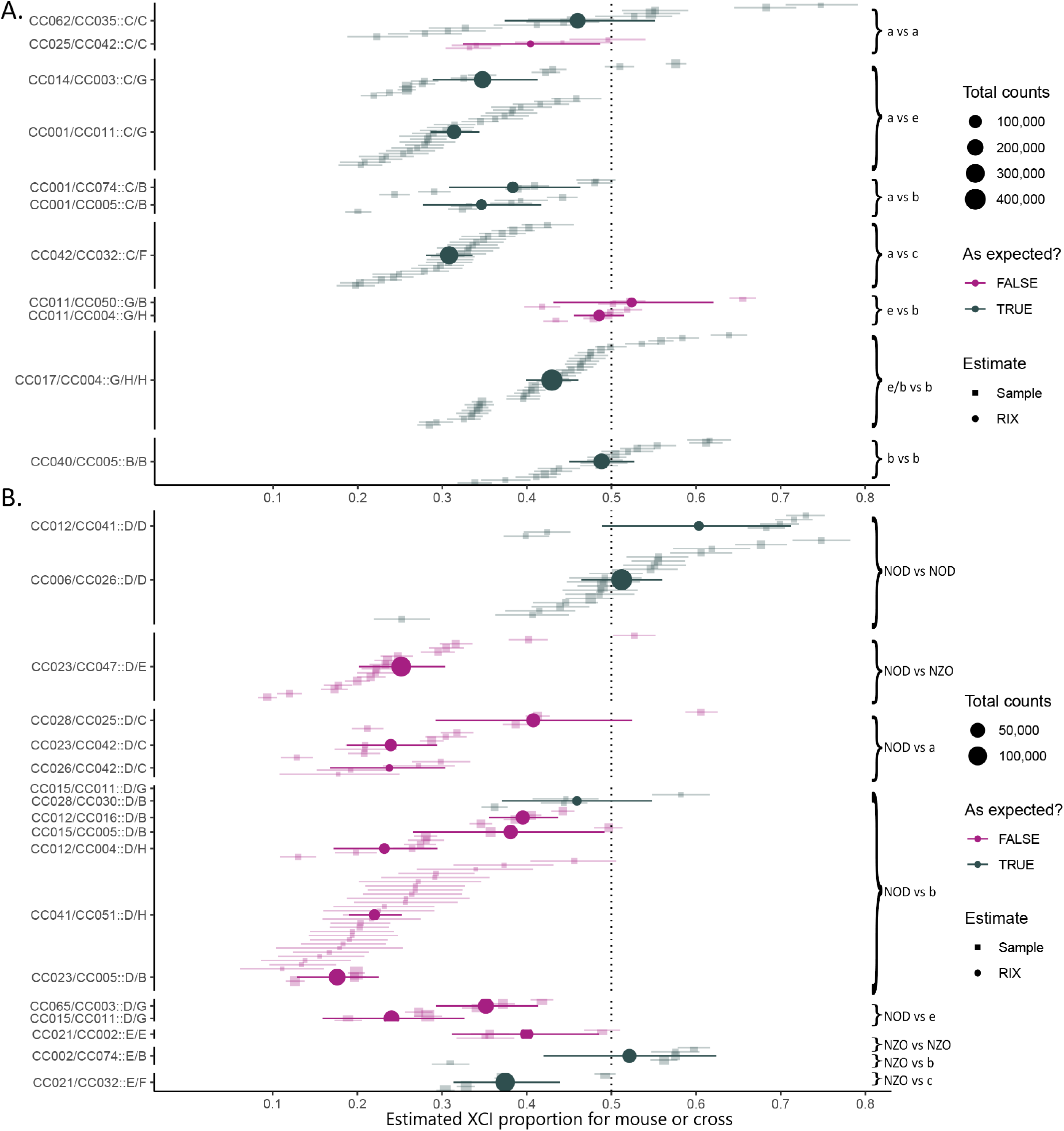
XCI proportion for all 266 samples across 28 CC-RIXs. Crosses where both CC parent contain previously-observed *Xce* alleles (A), and crosses where at least one CC parent is NOD or NZO within the interval (B). The y-axis labels state the CC-RIX followed by the two founder haplotypes that overlap the *Xce* in that CC-RIX. The *Xce* comparison for each group of RIX crosses is noted on the left vertical axis. Square points show the mean estimate of XCI proportion with 95% HPD bars for individual samples. The size of the point reflects the total k-mer counts from the sample, corresponding to its total RNA-seq read count and informativeness. Each cross is summarized across the RIX with round points. Crosses in gray match our predicted estimates of XCI skew based on known or inferred *Xce* allele whereas crosses highlighted in magenta do not.

Unlike the narrow HPD intervals seen at the gene and individual level (Figure 6), there is greater variability across individuals within a RIX. Some RIX from SP2 have wide HPD intervals reflecting their smaller replicate groups overall and perhaps a smaller starting amount of cells relative to SP1 due to RNA-seq sample collection for SP2 that took tissue from the striatum as opposed to whole brain tissue for SP1. An additional caveat is that haplotype reconstruction for SP2 relied upon genotyping data from the CC resource and not the specific individual mouse, which may have led to errors in assigning haplotypes, particularly near segregation points. Therefore, some RIX from SP2 have HPD intervals that are less informative, *e*.*g*. CC015/CC005, CC015/CC011, and CC021/CC002 (Figure 7).

The width of the HPD intervals at the RIX level derives from the precision, *α*_0_, of the overall RIX-wide XCI proportion. Though we described some legitimate experimental artifacts that may contribute to lower precision in certain RIX crosses, there are also true underlying biological reasons for this variation among samples in a cross. Inter-individual variability among the samples in a RIX can be interpreted as different amounts of starting cells that correspond to our precision estimate, *α*_0_, as described next.

### Estimated number of cells in pre-brain epiblast tissue range from 20 to 30

Our statistical model for sample-specific XCI proportion implies a Pólya-urn model for cell proliferation in which one of the estimated parameters, *α*_0_, relates to the number of brain precursor cells in the epiblast at the onset of random X inactivation. Our estimates of *α*_0_ were strikingly concordant between the two sample populations (Figure 8 and Table S2), and so we combined them to give a single, overall value. The combined posterior distribution for *α*_0_ followed a gamma distribution with shape parameter 100.36 and rate parameter 4.10. This translates to a point estimate (posterior mean) for *α*_0_ of 24.48 with standard error (posterior standard deviation) of 2.44 and a 95% HPD interval of 19.93 to 29.50. Our model thus suggests that the number of initial pluripotent cells in the epiblast that eventually form brain tissue in mature mice may be around 20-30. This is a reasonable figure given the number of total cells in the epiblast ranges from around 120 on E5.5 to 660 on E6.5 (Snow 1977).

**Figure 8.**
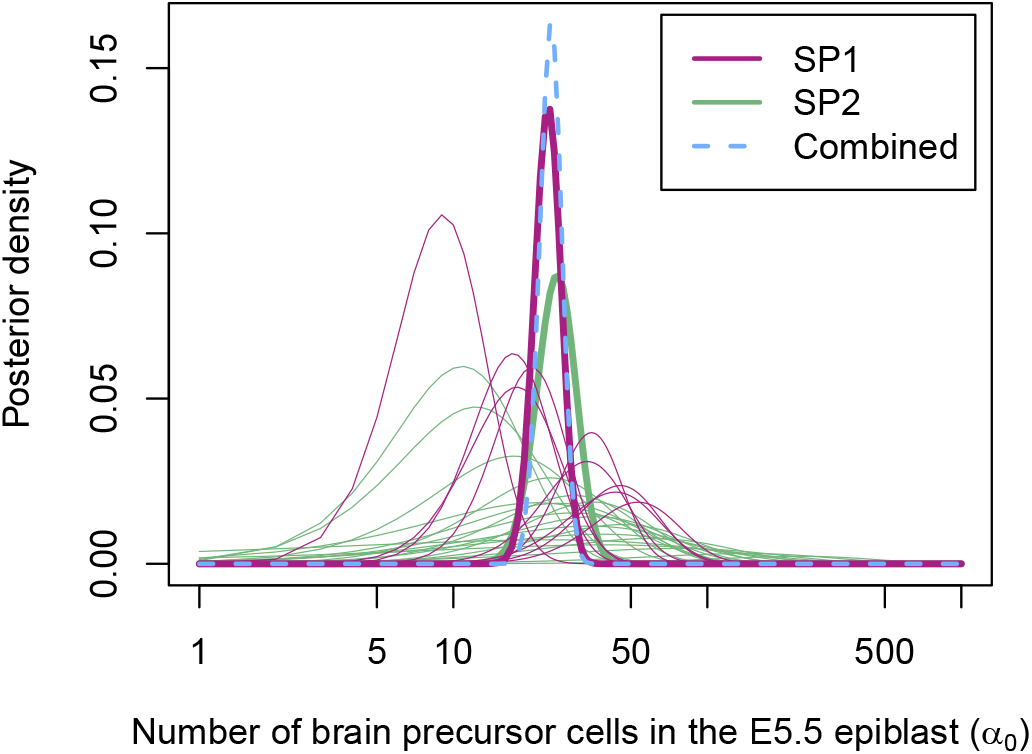
Bayesian inference of parameter *α*_0_, which estimates the number of brain precursor cells in the E5.5 epiblast. Fitted posterior curves are shown for each RIX (thin lines) from SP1 (blue) and SP2 (green), with consensus posteriors for SP1 and SP2 (thick blue and green), and an overall consensus posterior (dotted magenta) centered at 24.48 (95% HPD 19.93-29.50). The horizontal axis is given on the log scale for readability. Shape, rate, mean, and variance estimates from the posterior for each line are provided in Table S2.

### Unexpected XCI skewing in RIX females with the NOD Xce allele

YesAs well as corroborating earlier studies, the CC strain data also characterized the XCI (and thus *Xce* subtype) for two founder strains that had not been previously evaluated. Both founder strains, NOD and NZO, had been previously assigned to *Xce*^*b*^ due to sequence similarity with the reference genome.

We found a striking pattern of skewed XCI in crosses containing haplotypes derived from NOD at the *Xce* interval. Crosses heterozygous at this locus between NOD and any other founder exhibited profoundly skewed XCI proportions, despite the expectation that skewing would behave similarly with other strains carrying *Xce*^*b*^. Our results indicate that NOD harbors a novel *Xce* allele conferring a lower tendency to remain active, weaker even than *Xce*^*a*^. Figure 6 shows examples of gene- and sample-wide estimates of XCI proportion in three different CC-RIX crosses, each with NOD contributing the *Xce* region for at least one of its inherited X chromosomes.

Chromosomes bearing the *Xce* interval derived from NOD were consistently more likely to be inactivated than any other *Xce* allele (Figure 7b). This consistency suggests that this observed skewing is due to underlying variation that is inherent to the NOD *Xce* haplotype and not CC strain-specific factors, leading us to establish *Xce*^*f*^ from NOD as new allele in the functional series. Unexpected skews were observed in 11 out of 12 CC-RIX where one parental chromosome inherits the NOD *Xce* allele. This concordance was irrespective of different CC and founder strains carrying the *Xce*^*f*^, and transcended different *Xce* pairings, suggesting that this result is genuinely due to primary skewing.

Both our findings of 1) consistent skewing in crosses with previously-characterized alleles and 2) a new *Xce*^*f*^ functional allele in the NOD haplotype suggest that *Xce* lies in a minimum region from 102.46-105.56 Mb, consistent with the known *Xce* interval (see Figure 9). This is the only region on the X chromosome where crosses with *Xce*^*f*^ share a heterozygous NOD haplotype. The cross CC062/CC035 F1 females delineate the lower bound of the range because both CC strains are predicted to be derived from 129S1 until 102.46 Mb. Similarly, both parental strains of CC041/CC012 are predicted to contain the NOD haplotype until 105.56 Mb. This region overlaps the Chadwick *et al*. (2006) and Calaway *et al*. (2013) intervals on which we had based our initial *Xce* assignment, thus confirming the importance of the locus. Interestingly, chromosomes from NZO behave like they carry *Xce*^*b*^ which follows our *a priori* assumptions. This narrows our focus of inquiry because both NZO and NOD are identical-by-descent in this region and harbor few SNPs compared with the mouse reference genome. As a result, we investigated whether 1) the observed XCI skewing phenomenon in NOD—and by extension, other *Xce* functional alleles—may be driven by chromosomal rearrangements and not necessarily sequence variation; or 2), NOD and/or NZO were improperly categorized as *Xce*^*b*^ based on haplotype similarity.

**Figure 9.**
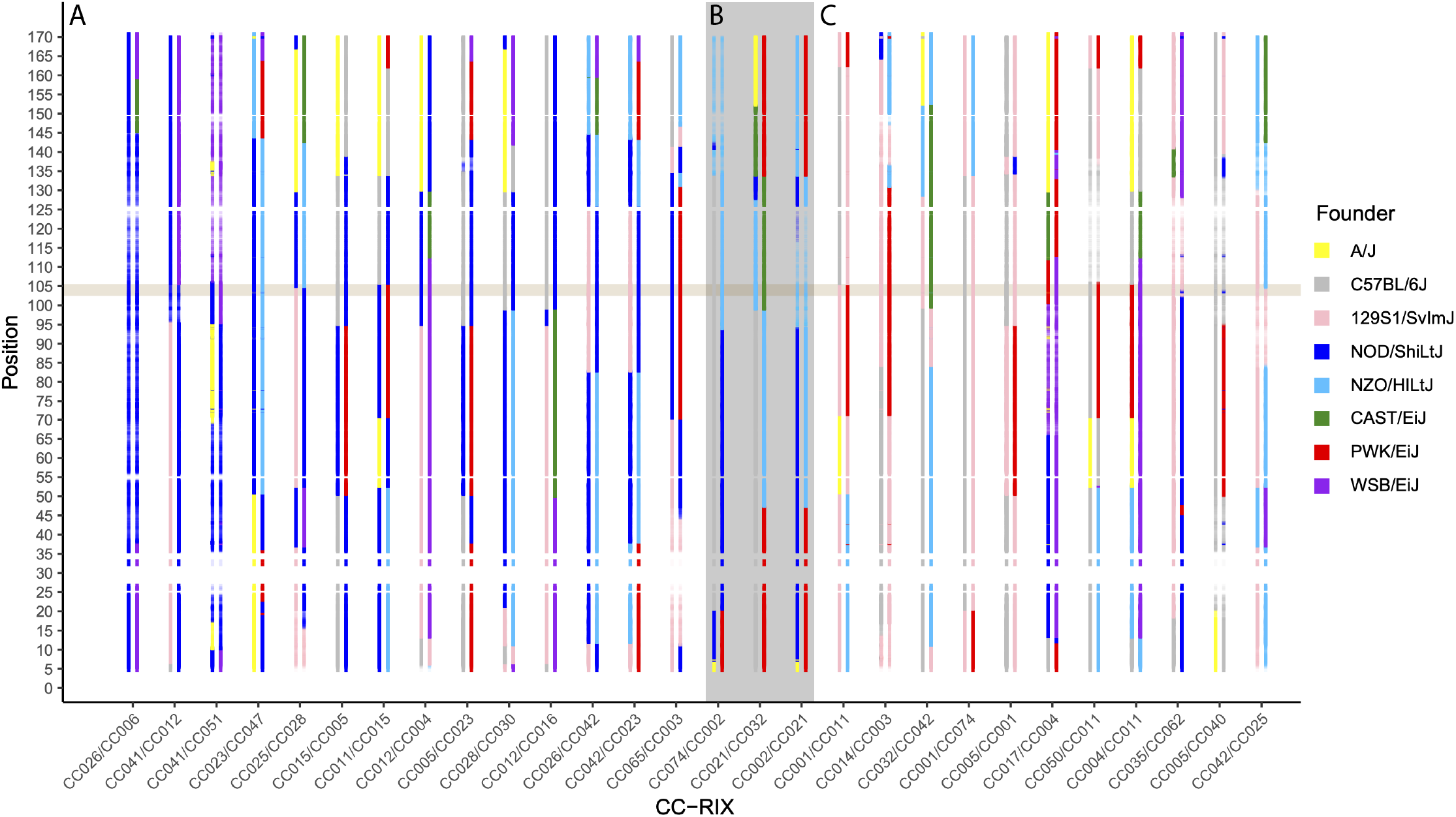
Heterozygous regions in each CC-RIX line illustrated by the predicted haplotype for each line. Haplotype assignments and their probabilities are represented by the color corresponding to each CC founder and the transparency of the colors, respectively. (A) The 14 crosses in this panel each contain the proposed *Xce*^*f*^. The beige highlighted region between 102.5 and 105.6 Mb is consistent with heterozygous regions in these crosses where exactly one founder is shared, namely, NOD. (B) These 3 crosses each contain the *Xce* interval derived from the NZO haplotype. (C) The 11 crosses in this panel contain only previously-characterized *Xce* alleles. The highlighted *Xce* interval region is consistent with the expected allelic series across the 14 non-NOD crosses and established *Xce* intervals.

### Analysis of SNPs in the Xce interval show that NOD and B6 have almost identical haplotypes

Given the unexpected patterns of XCI skewing in RIX females that carry the NOD *Xce* haplotype in heterozygosity, we decided to use recently released WGS from 75 CC strains to determine the extent of haplotype sharing between the CC founders. To ensure that we only compare orthologous sequences we limited this analysis to genomic regions spanning the *Xce* candidate interval that have copy number one in the reference genome and B6, and likely copy number one in each of the other CC founders. For each region, we assembled the CC founder sequence using the CC strain with the corresponding haplotype and deepest sequence coverage. After aligning each region, we used standard phylogenetic analysis to determine the relationships between the founder haplotypes (Figure 10). The results were fully consistent with the previously published haplotype sharing based on microarray genotyping (Calaway *et al*. 2013). Briefly, the eight founders are distributed in four well supported haplotypes: one represented by CAST, the second by PWK, a third that includes 129S1 and A/J; and the fourth and last comprises B6, NOD, NZO and WSB. We conclude that the expectation that NOD should be *Xce*^*b*^ is supported by haplotype sharing.

**Figure 10.**
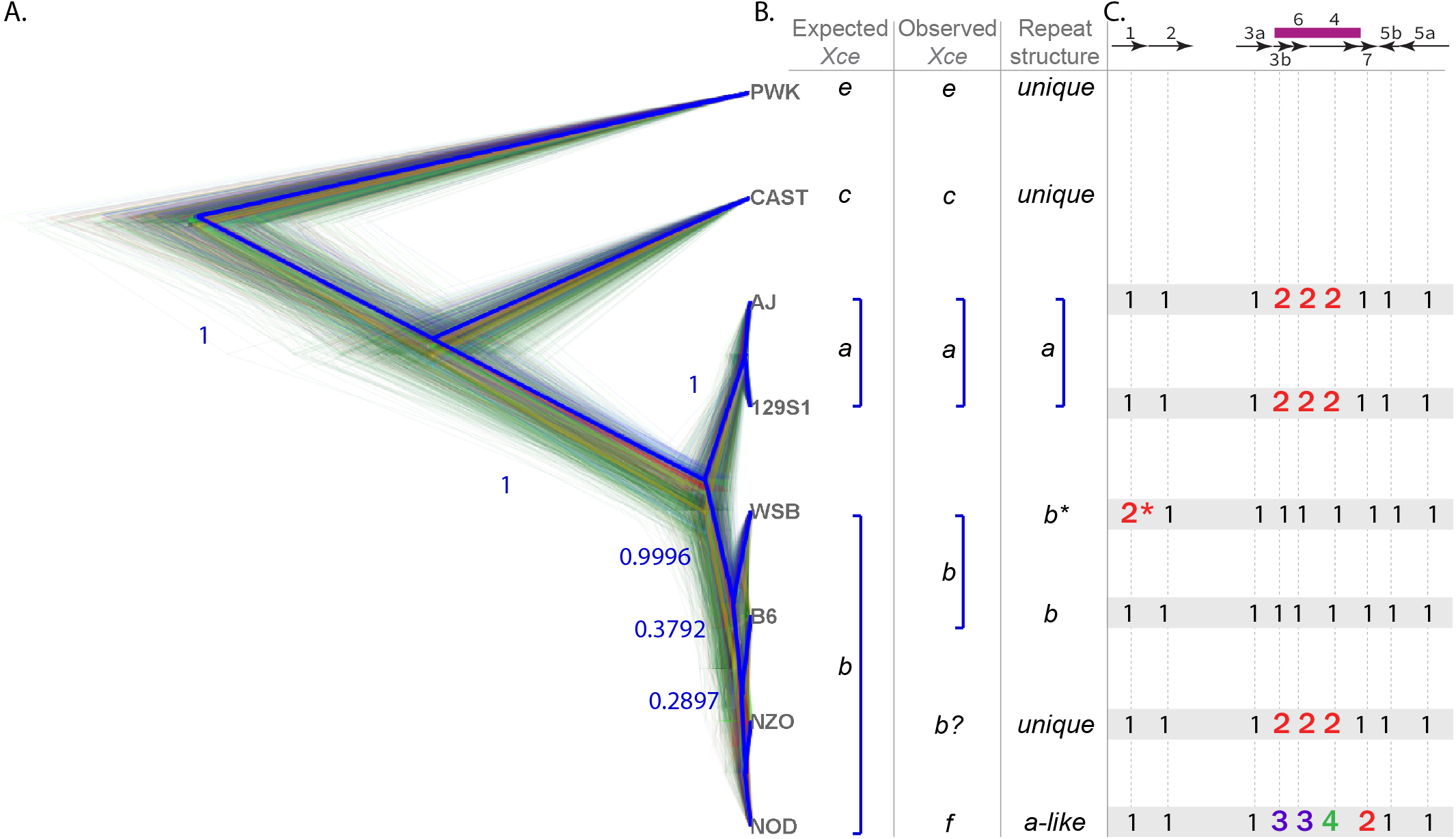
Phylogenetic tree generated using 8 sequences from the *Xce* interval among the 8 CC founder strains. A) Trees sampled from a Bayesian posterior of phylogenies, with the most frequently occurring topologies in blue, followed by green and red, respectively. The maximum clade credibility tree is shown in a thick blue line and posterior probabilities for each node in this consensus tree are shown next to the branch break point. B) Comparison of the expected *Xce* functional alleles based on haplotype similiarity for each of the founder strain, along with observed *Xce* strength and CNV repeat structure. C) Table of the observed number of copies for each segmental duplications (SD) and inversions (I) in the *Xce* interval. *The CNV landscape of WSB is similar to that of B6 except for a small duplication at the proximal end of SD1, which does not appear to affect XCI skewing. The SD and I pattern follow that described by Calaway *et al*. (2013) and Sheedy (2012), expanded to allow for more complicated duplication structures observed across the CC founder strains.

### Copy number variations distinguish weaker Xce alleles from stronger ones

The minimum *Xce* interval between 102,747,920-102,924,411 bp identified in Calaway *et al*. (2013) contain a series of recurring chromosomal rearrangements. These CNVs—segmental duplications (SD) and inversions (I)—were also verified in B6 with molecular assays by Sheedy (2012). We further corroborated the chromosomal architecture of this region in the mouse reference sequence with local nucleotide comparisons (Figure 11a) and optical mapping data (Figure S2). These rearrangements (detailed in File S10) have been posited as a potential explanation for the effect of the *Xce* functional allele series.

**Figure 11.**
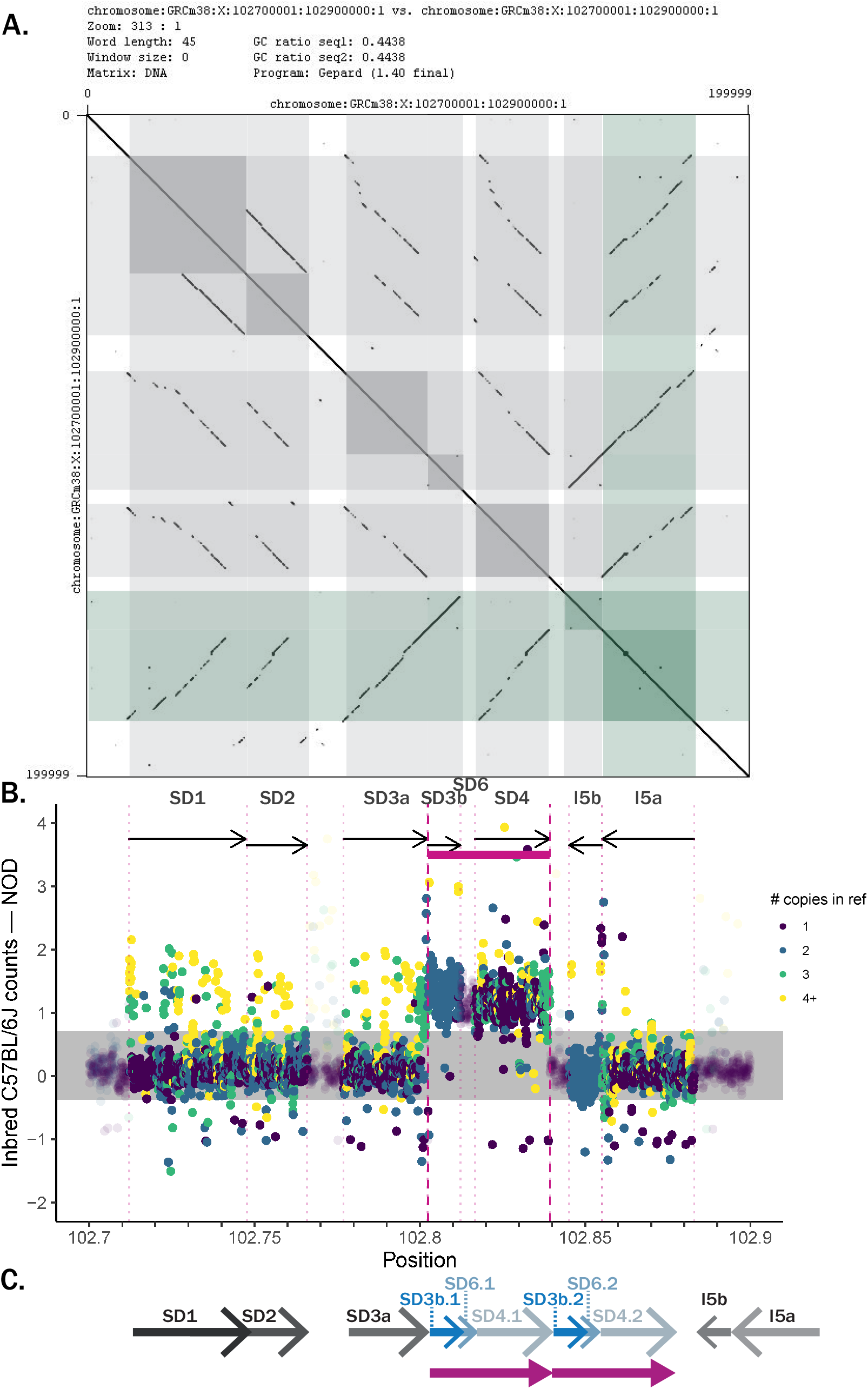
A) Dotplot of the mouse reference X chromosome from 102.7-102.9 Mb generated from pairwise sequence concordance in the genome assembly. Diagonal lines slanting down from left to right (shaded in gray) are duplications, while diagonal lines from left to right (shaded in green) are inversions. B) Difference in average counts of genomic 45-mers between sequenced samples with *Xce* haplotypes derived from NOD (16 CC strains and 1 inbred NOD) and counts from 24 inbred B6 strains. Arrows signify duplications (SD1-4) and inversions (I5a-b). There is a clear increase in copy number in the interval marked in magenta, R1. CNV clusters are centered at -0.895, 0.107 (shaded in gray), and 1.311. C) Schematic showing the hypothesized architecture of recurrent duplications and inversions within the *Xce*. The arrows in blue comprise R1, in magenta.

Using direct searches of non-overlapping 45-mers (File S5), we discovered an additional copy of the X chromosome sequence from approximately 102,802,400-102,839,400 bp, forming a continuous, 37 kb repeat spanning SD3b, SD4, and the bridge sequence between these recurring regions that we denote SD6. As shown in Figure 11b, the pertinent duplicated region marked with a magenta band clearly spans 45-mers with a consistent increase of counts centered at one extra copy. Henceforth we refer to this novel CNV as R1. Furthermore, we demonstrate that both A/J and 129S1, which both carry *Xce*^*a*^, share the same duplicated region, R1, as NOD with a roughly increased copy number of one (see Figures S4-S5). Replicated experiments over decades (Johnston and Cattanach 1981; Simmler *et al*. 1993; Calaway *et al*. 2013) have demonstrated that *Xce*^*a*^ was the weakest known *Xce* allele, previous to our finding in NOD.

This strong molecular evidence establishes a distinction between the reference genome and strains with weak *Xce* alleles, supporting the idea that variations in copy number within the *Xce* region contributes to the functional allele. We hypothesize R1 is associated with a weak *Xce* allele, and that the chromosomal organization of CNVs in NOD, A/J, and 129S1 may be described with the schematic shown in Figure 11C.

Compared with NOD, both A/J and 129S1 have hundreds of nucleotide variations relative to the reference (Figures S4-S5). Although all three strains share a similar pattern of repeats with R1, NOD has a weaker phenotype still compared with *Xce*^*a*^. Allelic series require multiple causal variants within the same locus and we hypothesize that additional variants between these two weak functional alleles explain the differences beyond their shared, similar CNV. Both XCI skewing and genetic differences still remain between NOD and the two strains confirmed to possess *Xce*^*a*^, leading us to establish NOD as its own allele in the functional series, *Xce*^*f*^.

Strikingly, the CNV pattern seen in NZO contains notable departures from those in other strains. NZO appears to have a more complex series of nested repeats such that different portions of the “weak repeat,” *i*.*e*. R1, are replicated at different frequencies (Figure 12). It carries three additional copies of SD4, two additional copies of SD3b, and one additional copy of a sequence segment distal to SD4 that we denote SD7.

**Figure 12.**
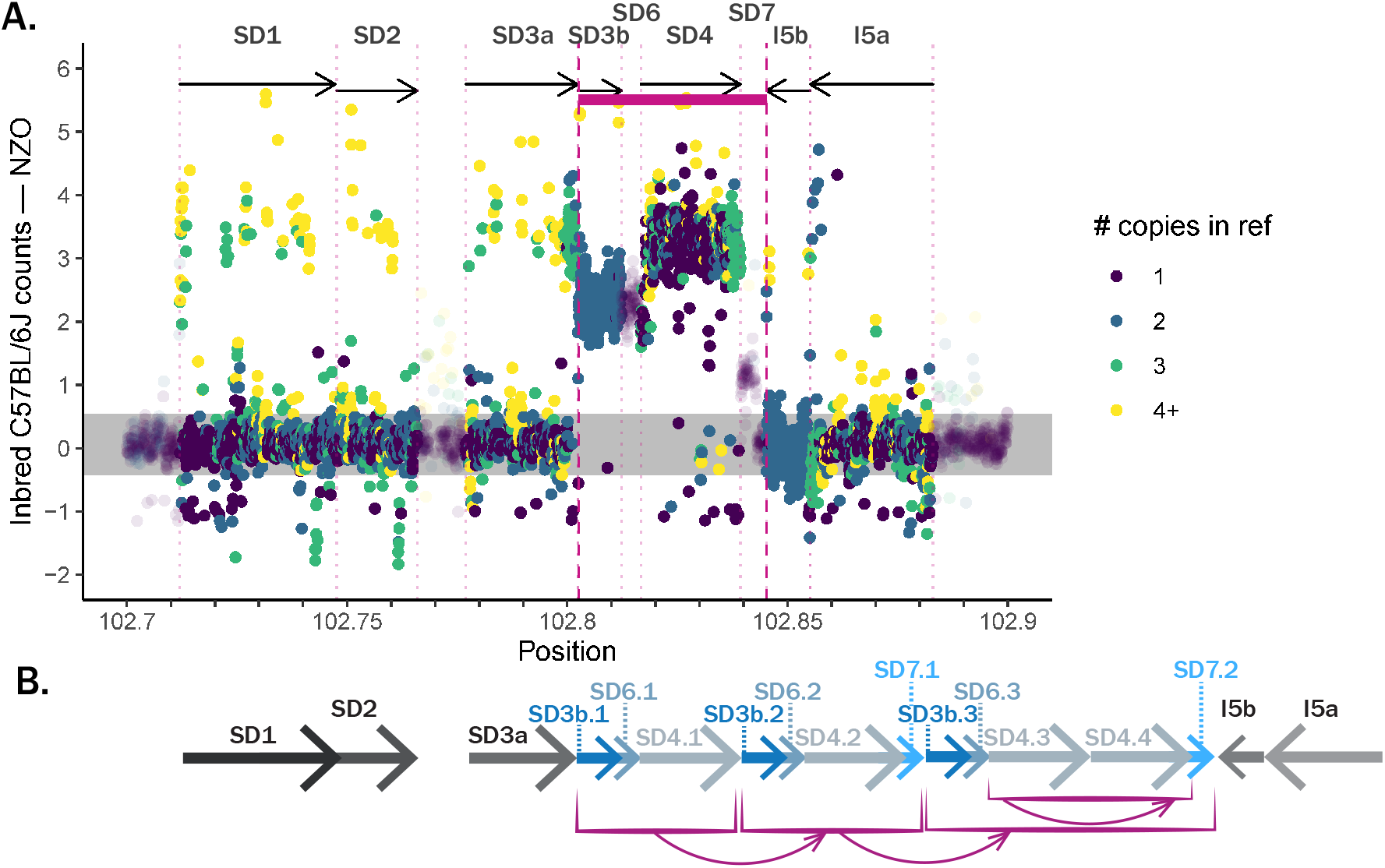
A) Difference in average counts of genomic 45-mers between sequenced samples with *Xce* haplotypes derived from NZO (7 CC strains and 1 inbred NZO) and counts from 24 inbred B6 strains. CNV clusters are centered at -0.878, 0.144 (shaded in gray), and 2.947. There are 62 SNPs across the 20 kb interval, 10 of which are in R1 (0.013% of the k-mers in the interval). B) Schematic showing the hypothesized architecture of recurrent duplications and inversions within the *Xce*.

We confirm that NZO has unique breakpoints between SDs that NOD and the reference sequence lack by querying matches of 45-mers at the SD boundaries. Neither the reference nor NOD contain repeats of SD7, so there is only one set of sequences flanking both sides of SD7, *i*.*e*. between SD4-SD7 and SD7-I5b. NZO, on the other hand, contains two distinct sets of k-mers on both the proximal and distal ends of SD7 (see Figure S9). This provides evidence that there are two copies of SD7 in NZO, one of which is a repeat flanked by sequences that form a pattern neither observed in the reference nor NOD. Although we are not able to verify the exact locations and pattern of the NZO duplications, shown as a hypothesized schematic in Figure S9B, we do see molecular evidence supporting the quantity of repeats in NZO and the presence of unique breakpoints between duplications and inversions. This suggests that NZO has a different chromosomal architecture in this region compared with other strains, though one that does not manifest in differences of XCI pattern compared to the reference strain.

## Discussion

In a previous study by our group (Calaway *et al*. 2013), we used a diverse set of inbred strains and allele-specific gene expression to characterize a new *Xce* phenotype and to narrow the historically well-defined *Xce* interval. That study identified a set of recurrent duplications within the *Xce* and suggested that variation in their copy numbers may in fact be the functional variation driving the allelic series. In the present study, we examined that hypothesis and quantified the skewing phenotypes of two CC founder strains with inferred *Xce* alleles based on sequence similarity with B6 across the *Xce* locus.

### Leveraging increased genetic diversity in CC-RIX identifies novel XCI patterns

Two important features of our methods are worth noting: 1) increased heterogeneity in the genetic composition of our F1 crosses of well-described CC strains, and 2) improved mapping resolution across the X chromosome from a novel method of quantifying ASE in CC-RIX mice and modeling the resulting counts in a hierarchical Bayesian manner. The animals represented in our study are each mosaics of 8 inbred mouse strains, with one X chromosome inherited entirely from each parent. Haplotype estimates across the genome in the CC strains are stable and replicable, allowing us to leverage previously collected genotyping and sequencing data to inform ASE estimates in our dataset. The haplotype reconstruction across the X chromosome for every cross used in this study is depicted in Figure 9.

The observed XCI skewing suggests that the *Xce* lies in a minimum region from 102.46-105.56 Mb based on haplotype probabilities from the genotyped CC-RIX in our study set and previously genotyped CC strains (Figure 9). Even though we had incorporated prior information about the *Xce* region based on results from Calaway *et al*. (2013) and Chadwick *et al*. (2006), those assumptions have held up to our results because our presumed *Xce*^*f*^ crosses share no other region in common. In addition, the crosses with well-characterized *Xce* alleles lack NOD in that region and are consistent with their known *Xce* subtypes.

Our methods relied upon a novel way to quantify ASE across the X chromosome by querying a set of curated 25-mers among the RNA-seq reads from each of the 266 mice in our study population. The 25-mers specifically targeted reference and alternate alleles at known polymorphisms in coding regions, and fed into a hierarchical Bayesian model to quantify XCI proportion for each cross and sample. Among the *Xce* alleles that have previously been characterized, our estimated XCI proportions matched what we would expect based on data from the literature (see Figure 7a). This finding serves to corroborate historical observations and to provide validation for our *Xce* imputation method and statistical model.

We observed highly variable proportions in some crosses, potentially owing to multiple sources of variation. Some CC strains have segregating boundaries at or near the *Xce* interval, making the assignment of CC strain from which the haplotype derives more uncertain, such as near 102.5 Mb in CC062. As shown in Figure 9, CC062/CC035 defines the lower boundary of the maximum *Xce* interval because the data is consistent with the XCI ratio being 50:50, *i*.*e*., between two *Xce*^*a*^ functional alleles of equal strength. In reality, CC062 has a large recombination interval between 129S1 and NOD near this proximal boundary. The broad range of proportions we actually observe suggests that *Xce*^*a*^ /*Xce*^*a*^ may not be an appropriate designation for every sample in this cross and that some may indeed be *Xce*^*a*^ /*Xce*^*f*^.

In addition, we have few samples and crosses with *Xce*^*e*^ derived from PWK. Our findings suggest that it is similar in strength to *Xce*^*b*^, consistent with previous findings (Calaway *et al*. 2013). As noted in the Methods, samples from SP2 had fewer replicates, leading to estimates of XCI skew that are less certain and more susceptible to any unaccounted-for RIX-wide variability. For example, CC021/CC002 F1 females are homozygous for NZO but show a slight XCI skewing centered at 0.4 (95% HPD 0.3117-0.4852). We note, however, that this estimate is based on only three females.. Lastly, RNA-seq tissue collection for SP2 used the striatum as opposed to whole brain tissue, resulting in a smaller starting amount of cells relative to SP1. XCI proportions for individuals in SP2 thus had higher variance, leading to less stable estimates and larger HPD intervals.

### Pólya urn-based approximation to the number of cells in pre-brain epiblast tissue

Inter-individual variability in XCI skew among genetically-identical samples within a RIX cross can be partitioned into experimental and biological variation. Although the two cannot be easily disentangled, we surmise that the biological variation derives, in part, from the precision of the beta distributed parameter for each estimate of mouse-specific XCI proportion. At the point of inactivation choice, the cells in the epiblast are akin to balls in a Pólya urn. The Pólya urn describes a random process in which an intial number of red and blue balls undergo successive rounds of randomly assigned duplications; after infinite rounds, the final proportion of red vs blue balls is a random number whose variability is a function of the total starting number. Urns that start with a greater number of balls are more stable against random fluctuations in the proportions of the red to blue balls, and have proportions more closely gathered around the starting proportion; urns starting with a smaller number lead to a final proportion that is more variable.

Analogously, the urn represents a RIX and *α*_0_ represents the starting number of pluripotent cells that are involved in the decision to activate either the maternal or paternal chromosome at around E5.5 and will eventually form brain tissue (or whichever tissue undergoes an ASE assay) in the mature mouse. Though we first estimate *α*_0_ in each RIX individually, we assume that the parameter should be similar in each individual cross, given the stability of biology underlying the XCI process.

Though we are unable to verify this quantity of 20-30 pre-brain epiblast cells, it does seem reasonable given the total number of cells in the epiblast is between 120-660 at E5-6 (Snow 1977).

### Copy number of recurrent duplications may explain the weakness of Xce^a^ and novel Xce^f^, found in NOD

Both NOD and NZO were previously predicted to express the *Xce*^*b*^ functional allele based on haplotype similarities to B6. Our results do not support this conclusion in NOD. We characterize the *Xce* locus derived from NOD as a separate functional allele in the series, *Xce*^*f*^, because we find it to be consistently weaker than all other known *Xce* haplotypes. Crosses involving 6 CC strains (CC012, CC023, CC026, CC028, CC041, and CC065) that contain the NOD-derived *Xce* region corroborate the weakness of the novel *Xce*^*f*^ (Figure 7). This continuity leads us to conclude that chromosomes carrying the NOD *Xce* allele contain sequence-level variation in this interval, manifesting in primary inactivation bias against keeping that parental copy of the X chromosome active.

We confirm that both NOD and NZO share sequence similarity in the *Xce* interval with the reference genome (Figure 10) using haplotype assembly from CC WGS and phylogenetic analysis. As a result, we conclude that CNV structure may be the causative factor for this phenomenon. CNV analysis (Figure 11b) reveals a large interval in which the normalized counts for all k-mers are consistent with the presence of an extra copy in NOD. We tentatively conclude that the repeat, R1, represents a genuine copy number increase of a contiguous 37 kb-long segment in NOD. R1 includes the entire SD3b and SD4, as well as the bridge sequence connecting them that is not duplicated in the reference (Figure 11b). The novel R1 appears to be recent; the last duplication found in the reference genome is that of SD3a-b inverting and inserting distally to form I5a-b, and is demonstrated by the sequence similarity between these two sets of sequences in both k-mer identity over sliding windows and optical mapping data. This general rearrangement structure is similar between two weak alleles, *Xce*^*a*^ and *Xce*^*f*^. A/J and 129S1 express *Xce*^*a*^, and both strains share with NOD evidence of the same SD3a-b to I5a-b inversion alongside the novel repeat, R1.

### NZO expresses Xce^b^ despite complicated CNV organization

XCI estimates from crosses containing an *Xce* region derived from NZO do not deviate from our hypothesized ratios based on the strain carrying *Xce*^*b*^. Whether the XCI proportions seen in NZO indicate that its *Xce* interval is the same molecular species with genuinely identical function as *Xce*^*b*^, or if the two phenotypes have converged to appear similar is unclear. We would expect NZO to have a duplication structure akin to that of the reference mouse genome, or at least a different structure to that of NOD, A/J, and 129S1. Our analysis of NZO is hampered by the lack of CC-RIX in our data with NZO in the *Xce* region. One of our main study populations, SP1, was designed to maximize heterozygous loci between B6 and NOD, which explains the predominance of both strains in our downstream analysis. Nevertheless, the three RIX that contain NZO are consistent with the strain bearing *Xce*^*b*^ or at least a functional allele of the same strength.

In NZO, we find a more complex pattern of SD’s and I’s than seen in other strains. As shown in Figure 12, NZO appears to harbor one increased copy of SD7, two increased copies of SD3b and SD6, and three increased copies of SD4. We confirm the increased copy number of these elements by observing novel sets of boundaries between SD’s that are not present in B6, NOD, or other strains. For example, NZO contains two distinct sets of sequences on the distal end of SD7, suggesting that there are two real copies of the segment in the NZO sequence: one of which leads into I5b and is present in the B6 sequence, and the other of which is novel (see Figure S9).

Thus, the copy number pattern observed in NZO is indeed different than what we observe in NOD, A/J and 129S1, and the reference genome. NOD and NZO were predicted to share the same skewing phenotype as the reference based on sequence similarity at the SNP level. Our data demonstrates that the NOD *Xce* haplotype has a novel functional allele, distinct from NZO and any known *Xce* allele. CNVs can explain the difference between the functional *Xce* alleles present in NOD and B6 but they are not able to discriminate between NOD and strains with the *Xce*^*a*^ allele. This is not particularly surprising given that this simplified approach ignores the potential effect of variation outside of the recent NOD duplication and do not consider higher order factors associated with duplications such as location and orientation of the duplicated segment. There are hundreds of sequence variants within *Xce* that differentiate NOD and the *Xce*^*a*^ haplotypes, and may explain the skewing differences between them. The similar CNV pattern between NOD, A/J, and 129S1 results in a weak *Xce* allele, however this does not preclude other sequence variants within the interval from playing a part as well. With the inclusion of the *Xce*^*f*^ allele described here, the series now contains six distinct functional alleles. As more *Xce* alleles are described, so increases the need for multiple causal variants in the interval.

### CNV abundance and organization, along with sequence variation, may all play a role in Xce strength and XCI skewing

Our data implicates copy number changes as being important to XCI, but there are additional factors distinguishing NOD from strains in *Xce*^*a*^, as all of these strains appear to contain the novel R1. In addition, the duplication structure found in NZO is more complex than what we observe in other strains yet this does not translate to a detectably different phenotype compared with B6. This suggests that alternate recurrent duplication structures, each containing variations relative to the reference mouse genome, may present technically different *Xce* species that converge in similar phenotypes. This is supported by phylogenetic analysis showing that A/J and 129S1 are more similar to each other, while NOD and B6 evolved separately along another branch. The larger region surrounding the *Xce* contains other recurrent duplications and repeats, indicating that it is a potential “hotspot” of copy number changes (Sheedy 2012). Further work into the nature of the *Xce* may explore the patterns and inheritance of those rearrangements. Broader molecular characterization of the extent that CNV plays a role in enacting this control will be required to fully understand the function of *Xce*.

## Acknowledgements

This work was funded by a National Institute of Mental Health (NIMH) grant R01-MH100241 to WV and LMT, NIMH and National Human Genome Research Institute (NHGRI) grants (P50-MH090338, P50-HG006582, and U24-HG010100) to FPMV, and a National Institute of General Medical Sciences (NIGMS) grant R35-GM127000 to WV. KYS and DO received partial support from NIGMS training grant T32-GM067553. KYS also received partial support from the UNC-CH Caroline H. and Thomas S. Royster Fellowship. VZ is funded via National Institute of Environmental Health Sciences training grant T32-ES007018. MiniMUGA was developed under a service contract to FPMV and other investigators at UNC-CH from Neogen Inc., Lincoln, NE. None of the authors have a financial relationship with Neogen Inc. apart from the service contract listed above. The authors have no other conflict of interest to declare. Members of Leonard McMillan’s lab at UNC-CH, including Maya Najarian and Sebastian Sigmon, provided guidance with the WGS data and building k-mers. James Xenakis was instrumental in accessing data from SP2. We thank Greg Keele and other members of the Valdar laboratory for helpful discussions and thoughtful comments on the manuscript.

## Supporting information

**File S1** Documentation for supplemental files (DOCX).

**File S2** Demographic information on 266 mouse samples, and summary of results (CSV).

**File S3** Complete data on 7,957 25-mers used to quantify gene expression on X chromosome (CSV).

**File S4** Posterior mode, mean, median, and 95% highest posterior densities determined by the Bayesian hierarchical model for all covariates in GLM performed for SP1 and SP2 (CSV).

**File S5** Complete data on non-overlapping genomic 45-mers used to determine CNV in *Xce* interval and resulting counts (CSV).

**File S6** Counts of 25-mers centered at a reference allele from SP1 RNA-seq data (CSV).

**File S7** Counts of 25-mers centered at an alternate allele from SP1 RNA-seq data (CSV).

**File S8** All counts of 25-mers from SP2 RNA-seq data (zipped directory containing CSV files).

**File S9** Plots of individual XCI proportion estimates across X chromosome for all 266 samples in study, akin to Figure 6 (PDF).

**File S10**: Positions of seven CNVs (segmental duplications and inversions) in *Xce* (TXT).

**Figure S1** DAG showing in detail the parameters of the hierarchical model for XCI proportion at the gene-, individual mouse-, and RIX level. Estimates for the number of day 5 brain precursor cells (*α*_0_) across RIXs are then combined through a post-processing step.

**Figure S2** Bionano optical mapping alignment for the mouse reference sequence.

**Figures S3-S8** Counts of 45-mers spanning the *Xce* and the difference between average counts in the indicated strain across the interval and counts from inbred B6 strains. The following strains are represented, respectively: B6, A/J, 129S1, CAST, PWK, WSB.

**Figure S9** Counts of k-mers spanning the proximal and distal ends of SD7 for both B6 and NZO. Sequence data from NZO clearly show at least two unique boundaries, suggesting that SD7 is indeed duplicated in this strain.

## Supplemental Figures and Tables

The following pages provide supplemental figures and tables for the manuscript *“Bayesian modeling of skewed X inactivation in genetically diverse mice identifies a novel Xce allele associated with copy number changes”* by Sun et al.

**Figure S1.**
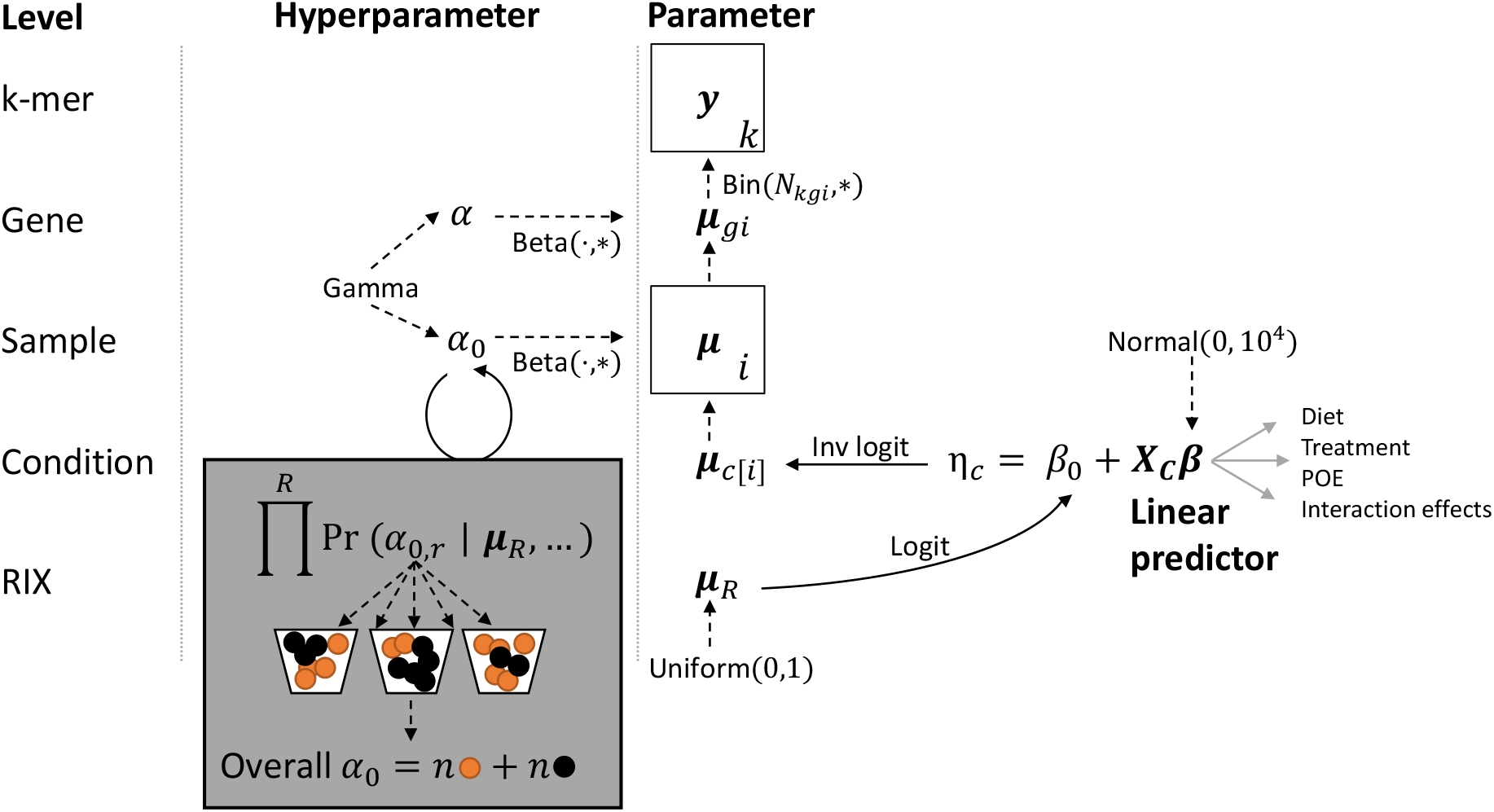
DAG showing in detail the parameters of the hierarchical model for XCI proportion at the gene-, individual mouse-, and RIX level. Estimates for the number of day 5 brain precursor cells (*α*_0_) across RIXs are then combined through a post-processing step. Arrows indicate dependencies between nodes. Dashed arrows are probabilistic dependencies, and labels denote the probability distributions linking the nodes. For distributions with multiple parameters, the star (*) indicates the parameter of the parent node, and the dot (·) is a placeholder for the other parameter. We use a parameterization of the beta distribution based on mean and precision, as described in the text. Solid arrows are deterministic dependencies, and labels denote the operation linking the nodes.

**Figure S2.**
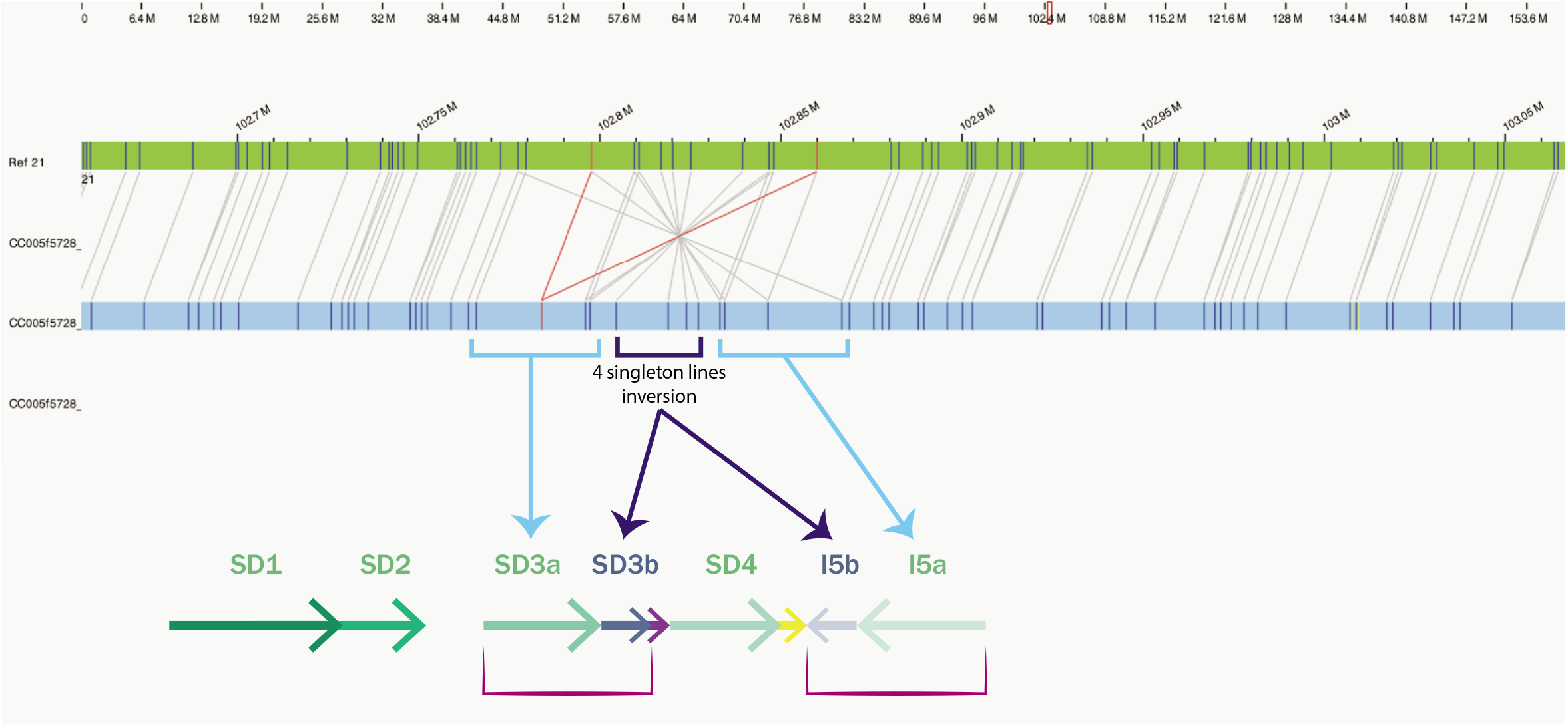
Bionano optical mapping alignment of the mouse reference sequence (labeled Ref 21) and CC005, which has a C57BL/6J-derived *Xce* region. Lines connecting the two sequences represent shared markers, and the red line linking one marker on CC005 and two on Ref 21 indicates that the marker on CC005 shows up twice in the reference. The shared lines in the marked region line up with the known duplication and inversion structure in C57BL/6J (blue arrows), and suggests that SD3a-b inverts and duplicates to form I5a-b.

**Figure S3.**
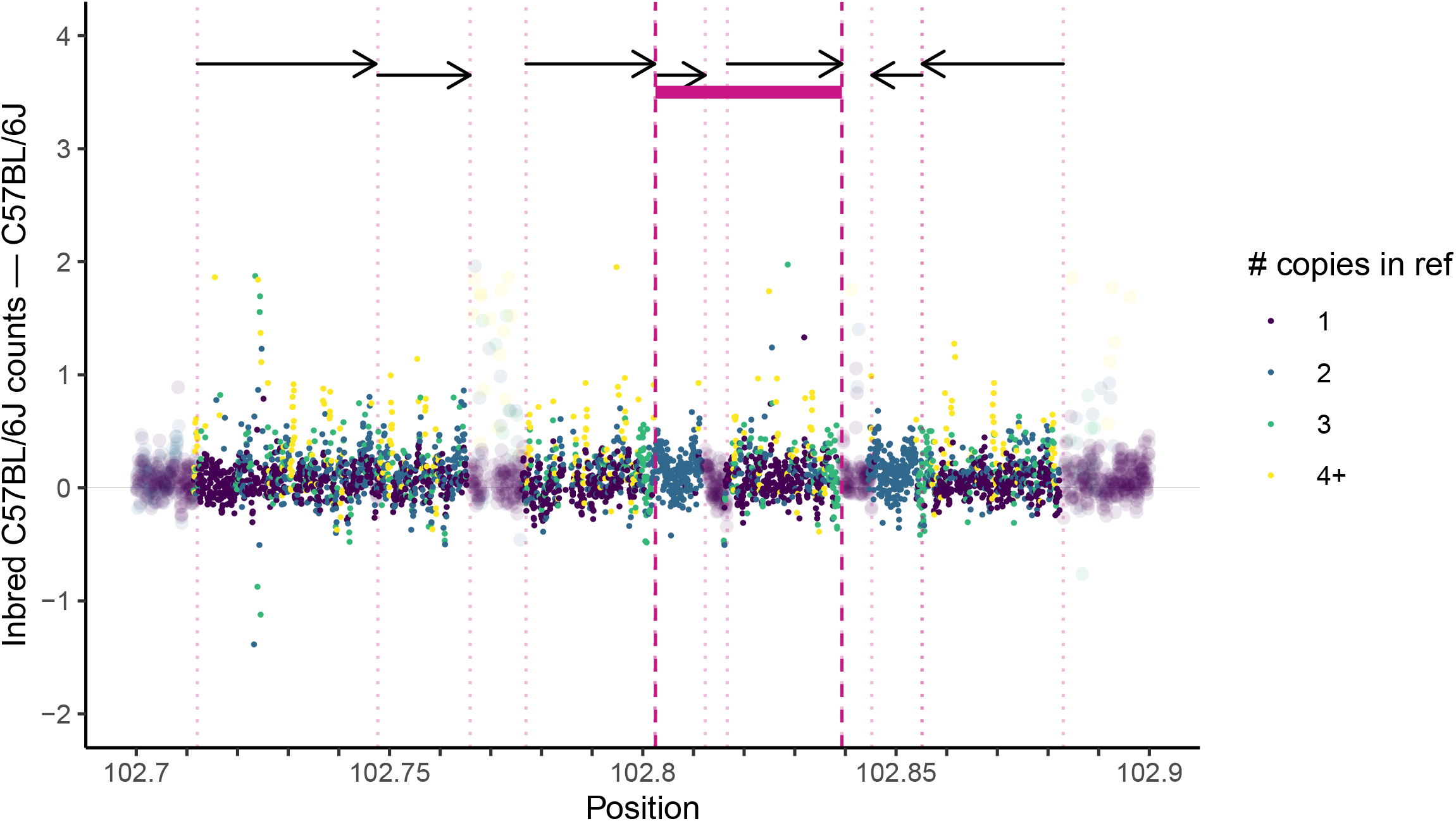
Counts of genomic 45-mers spanning the proposed *Xce*: difference between average counts across the interval in sequenced mice with haplotypes derived from C57BL/6J (10 CC strains) and counts from 24 inbred C57BL/6J mice.

**Figure S4.**
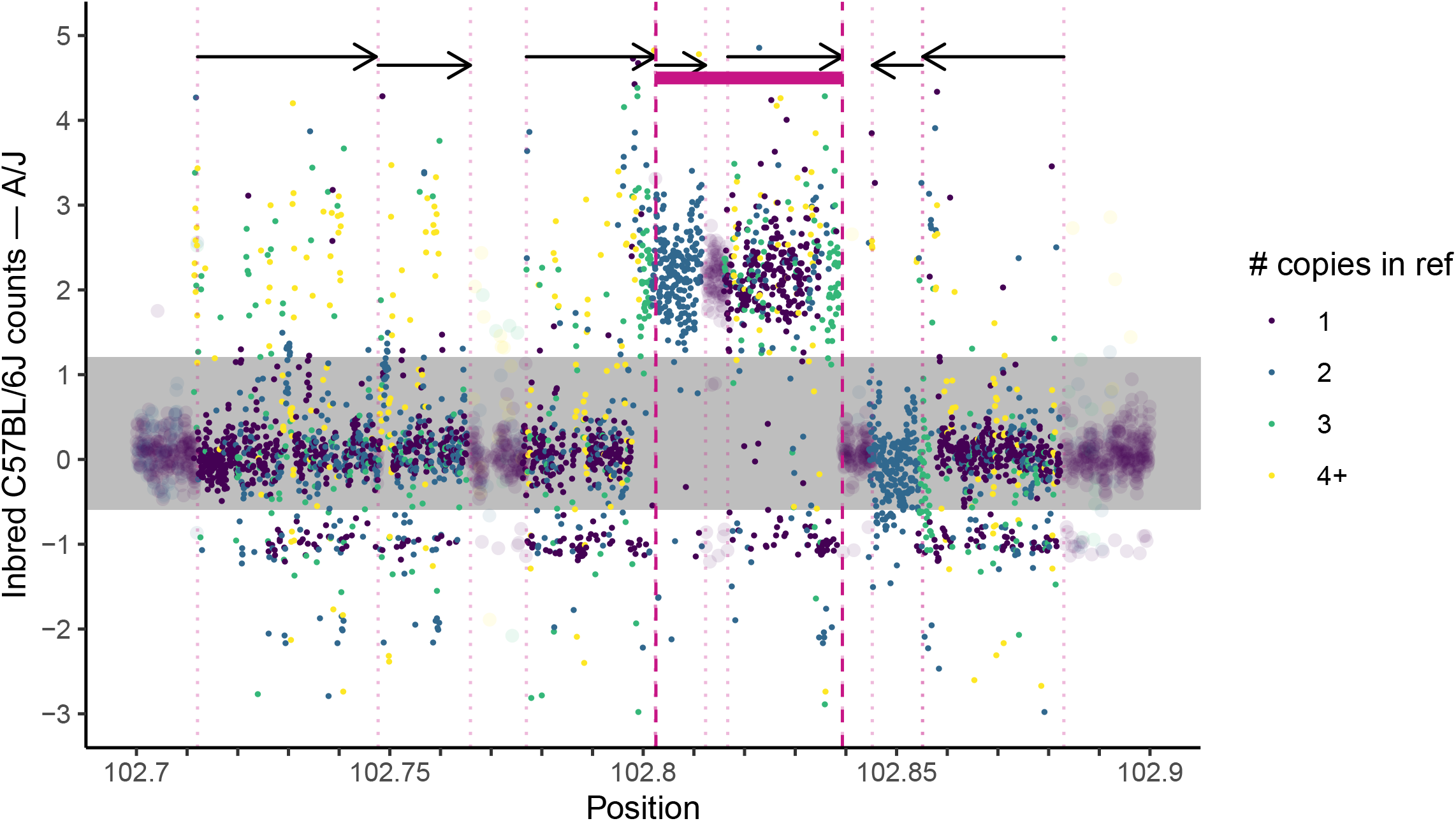
Counts of genomic 45-mers spanning the proposed *Xce*: difference between average counts across the interval in sequenced mice with haplotypes derived from A/J (4 CC strains and one inbred A/J representative) and counts from 24 inbred C57BL/6J mice.

**Figure S5.**
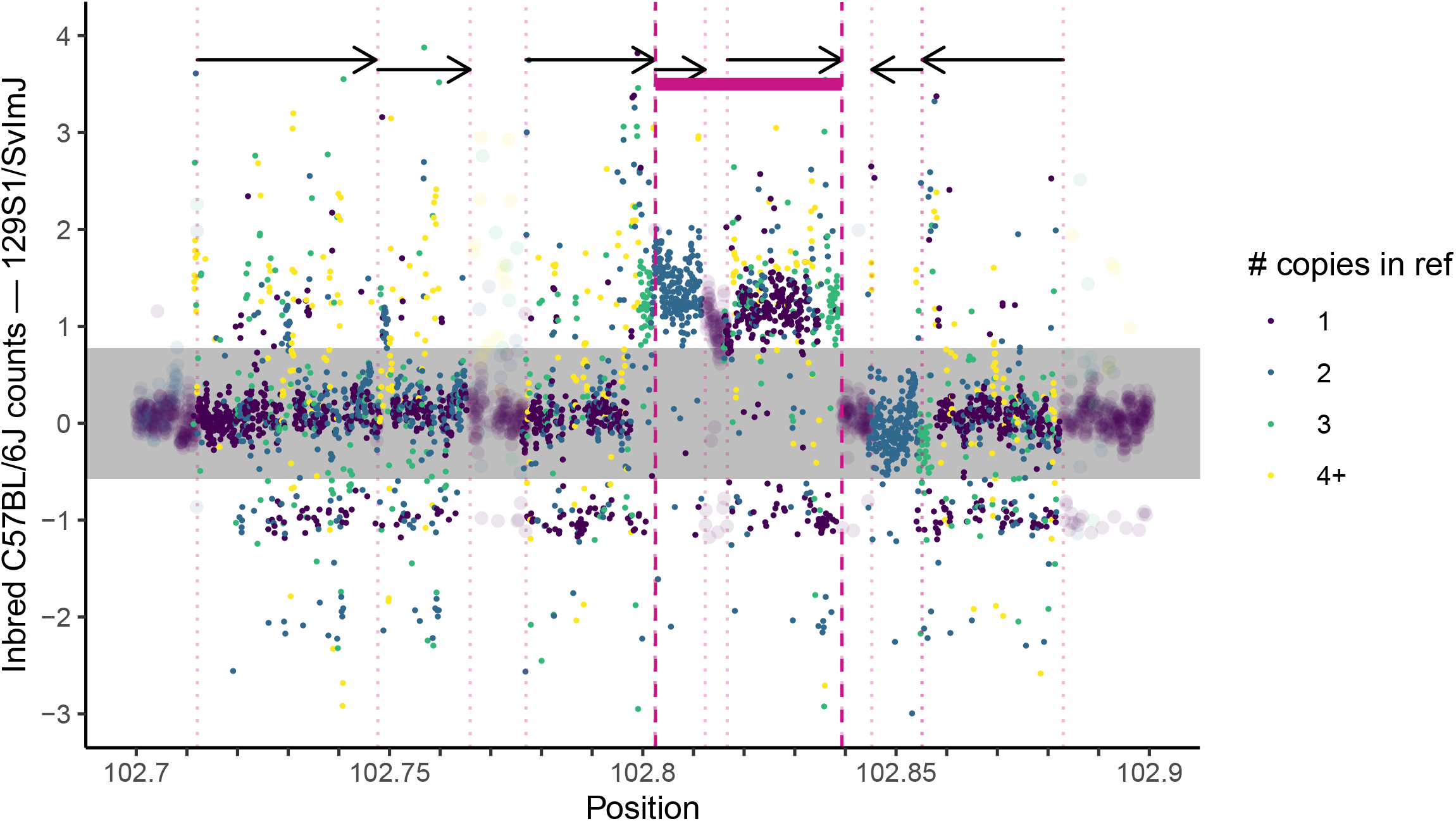
Counts of genomic 45-mers spanning the proposed *Xce*: difference between average counts across the interval in sequenced mice with haplotypes derived from 129S1/SvlmJ (15 CC strains and one inbred 129S1/SvlmJ representative) and counts from 24 inbred C57BL/6J mice.

**Figure S6.**
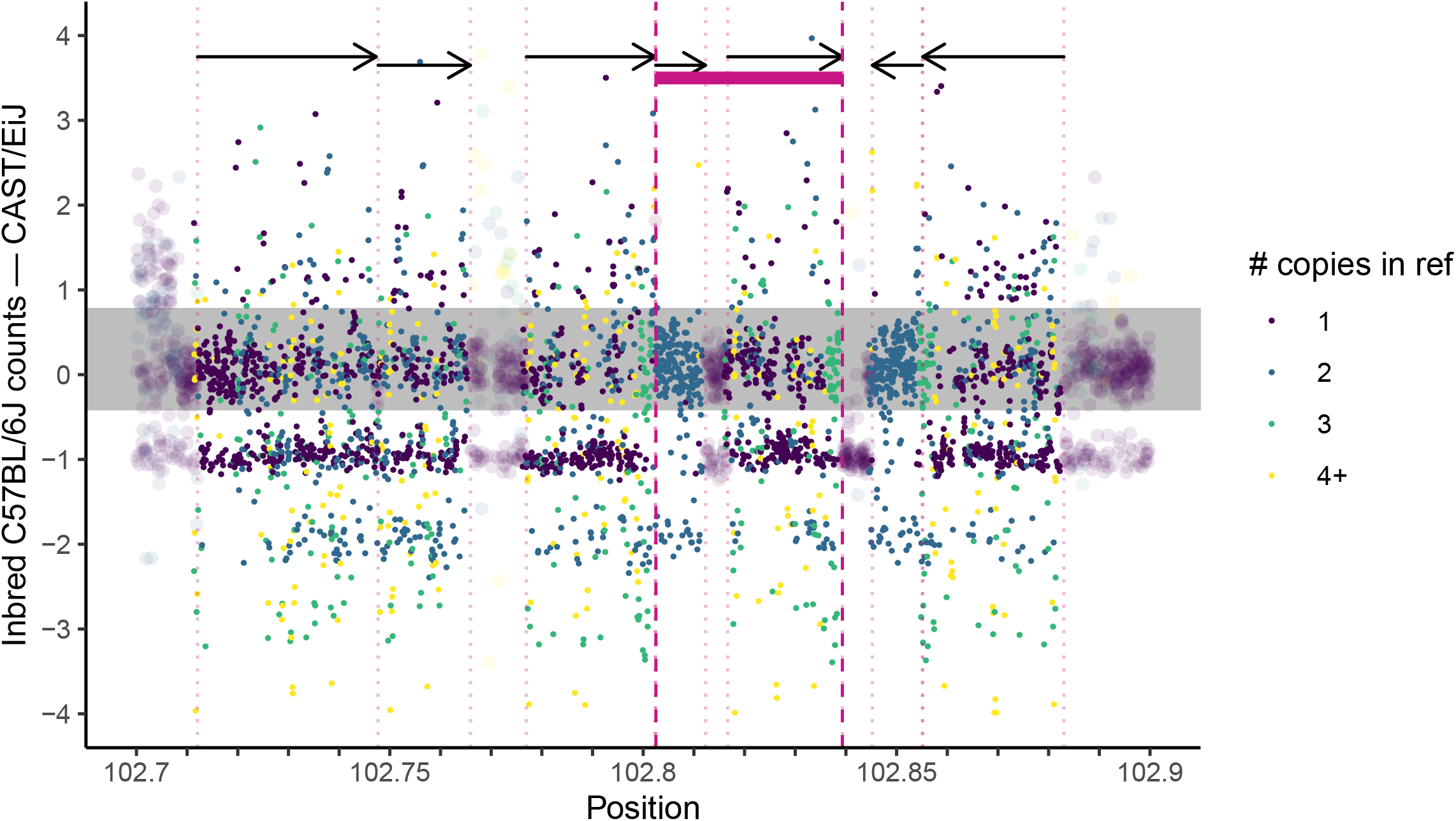
Counts of genomic 45-mers spanning the proposed *Xce*: difference between average counts across the interval in sequenced mice with haplotypes derived from CAST/EiJ (3 CC strains and one inbred CAST/EiJ representative) and counts from 24 inbred C57BL/6J mice.

**Figure S7.**
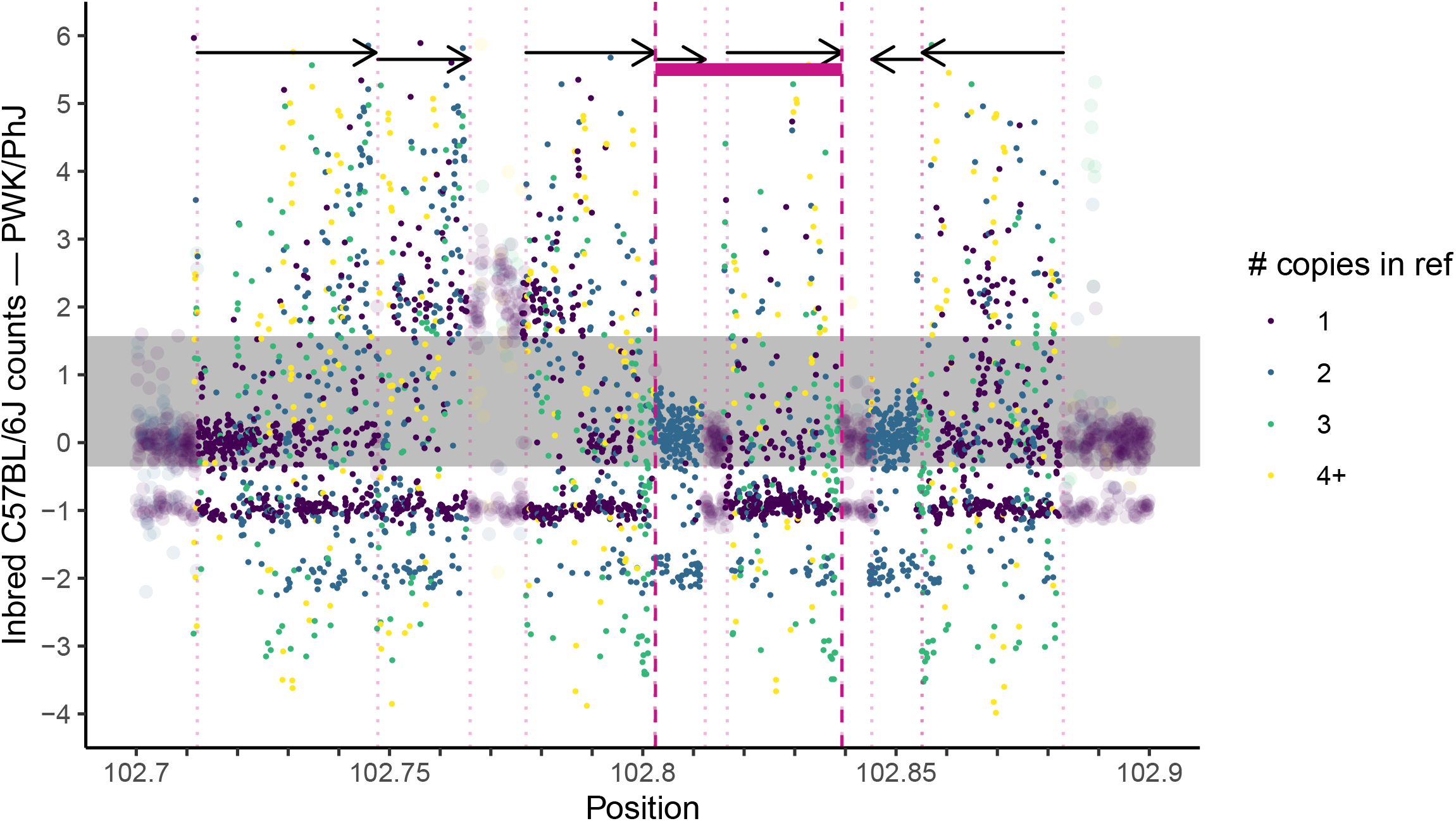
Counts of genomic 45-mers spanning the proposed *Xce*: difference between average counts across the interval in sequenced mice with haplotypes derived from PWK/PhJ (2 CC strains and one inbred PWK/PhJ representative) and counts from 24 inbred C57BL/6J mice.

**Figure S8.**
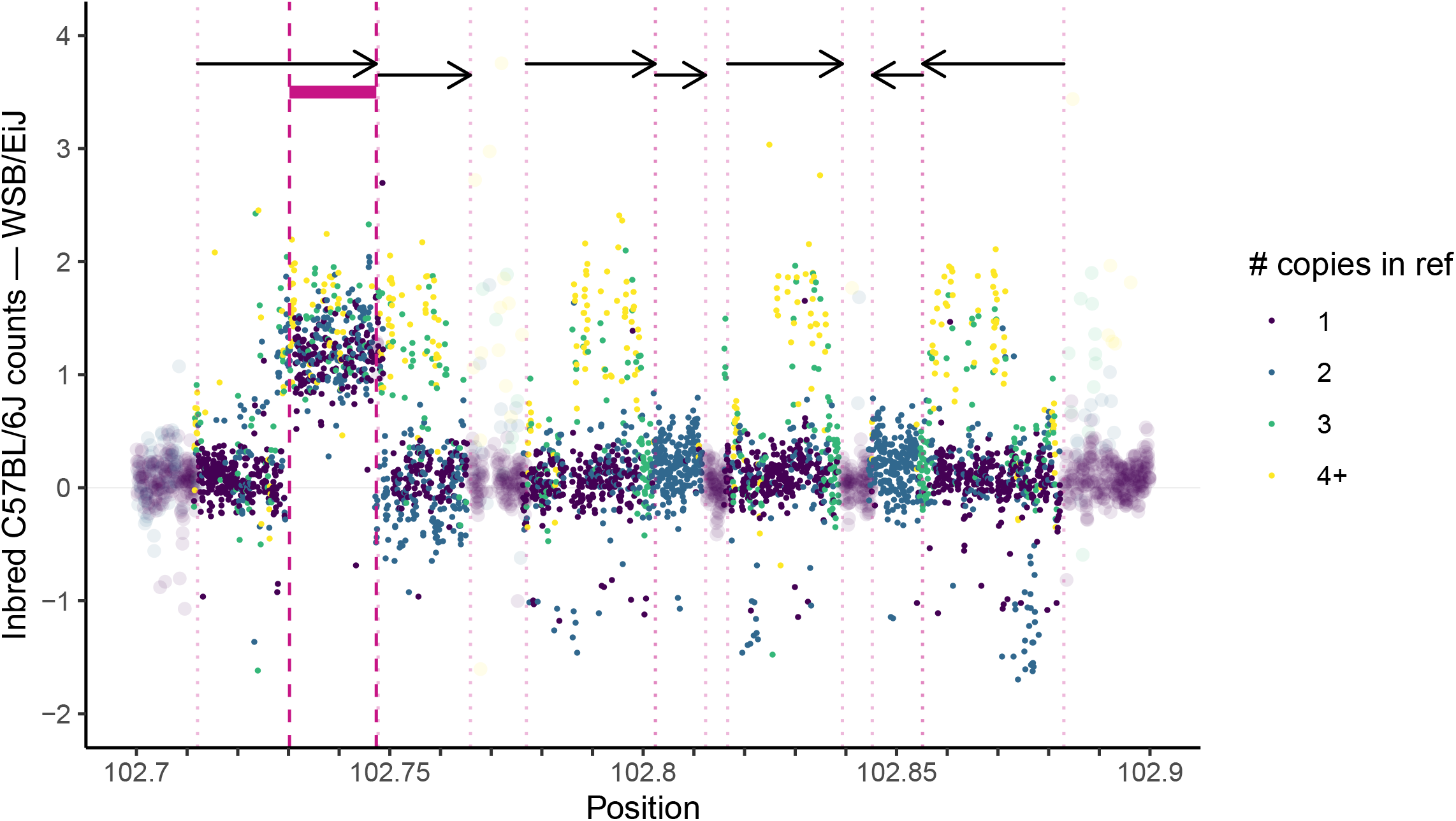
Counts of genomic 45-mers spanning the proposed *Xce*: difference between average counts across the interval in sequenced mice with haplotypes derived from WSB/EiJ (7 CC strains and one inbred WSB/EiJ representative) and counts from 24 inbred C57BL/6J mice.

**Figure S9.**
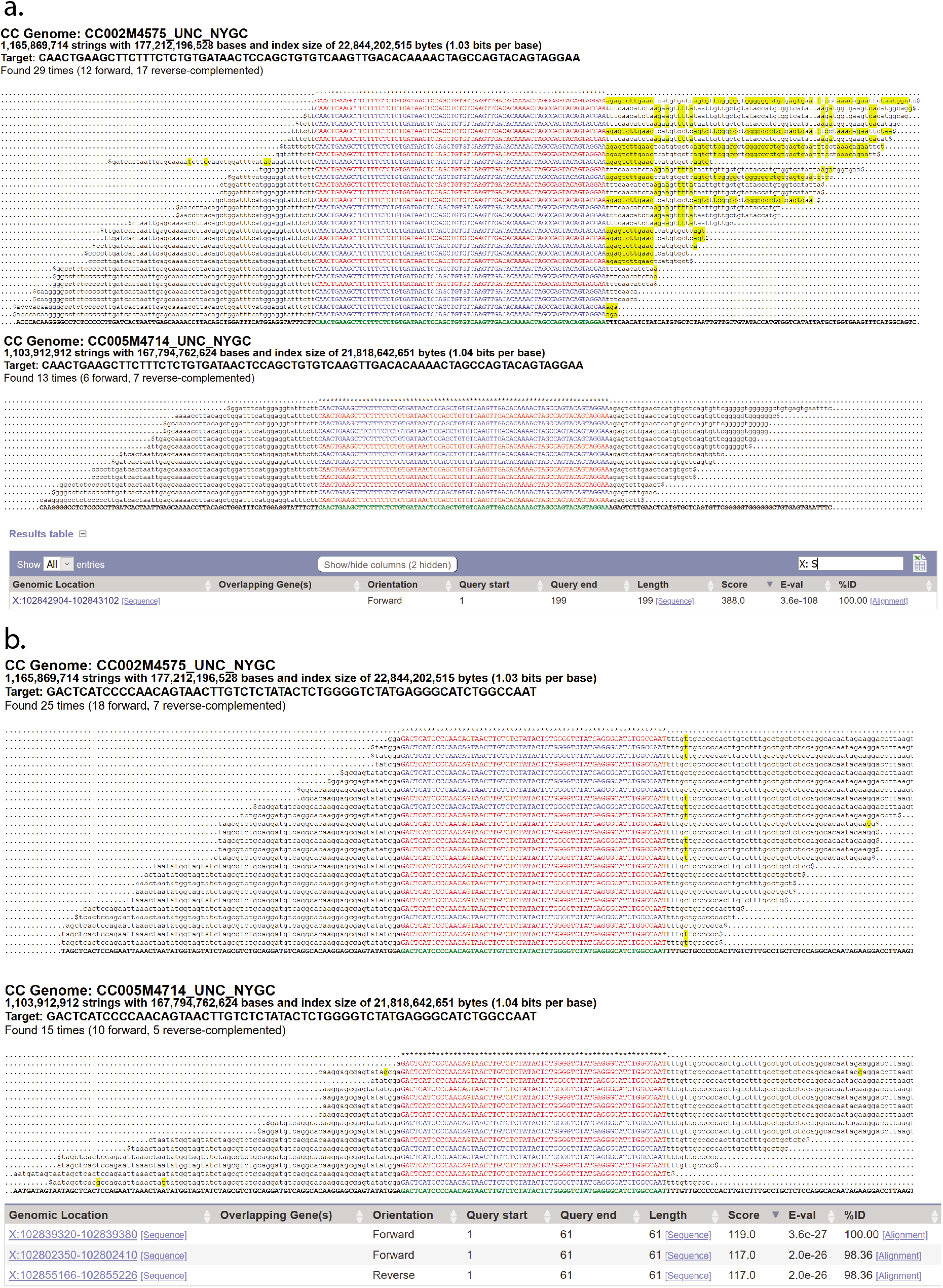
Counts of k-mers spanning the proximal and distal ends of SD7 for both C57BL/6J and NZO. Sequence data from NZO clearly show at least two unique boundaries, suggesting that SD7 is indeed duplicated in this strain.

**Table S1.**
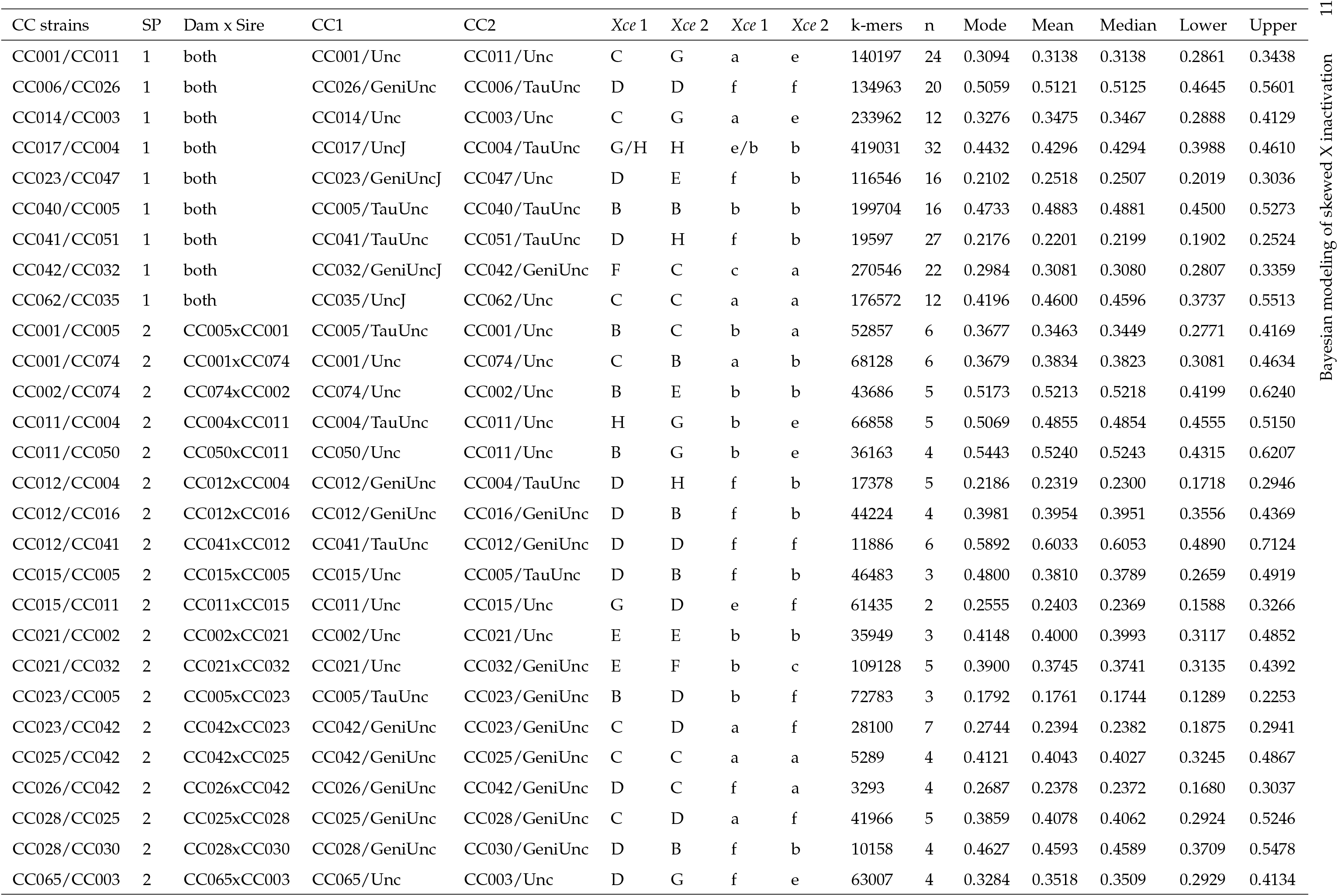
Summary of all RIX crosses phenotyped in this study, the number of samples per cross, the CC haplotype imputed in the *Xce* interval (first *Xce* 1-2 column), and the corresponding *Xce* alleles (second *Xce* 1-2 column).

**Table S2.** Shape, rate, mean, variance, and 95% HPD estimates for gamma-distributed *α*_0_ posterior distributions from MCMC run on individual CC-RIX.

| RIX | SP | Shape | Rate | Mean | Var | Lower | Upper |
| --- | --- | --- | --- | --- | --- | --- | --- |
| CC001/CC011 | 1 | 8.4312 | 0.1636 | 51.5364 | 315.0213 | 22.8404 | 91.7086 |
| CC006/CC026 | 1 | 10.3473 | 0.4592 | 22.5339 | 49.0736 | 10.9658 | 38.1977 |
| CC014/CC003 | 1 | 6.8406 | 0.3285 | 20.8238 | 63.3904 | 8.2642 | 39.0861 |
| CC017/CC004 | 1 | 13.2852 | 0.3512 | 37.8331 | 107.7394 | 20.3036 | 60.7270 |
| CC023/CC047 | 1 | 8.6800 | 0.4467 | 19.4306 | 43.4965 | 8.7338 | 34.3314 |
| CC040/CC005 | 1 | 6.7684 | 0.1324 | 51.1223 | 386.1305 | 20.1660 | 96.2263 |
| CC041/CC051 | 1 | 7.9745 | 0.2073 | 38.4634 | 185.5208 | 16.5788 | 69.3987 |
| CC042/CC032 | 1 | 7.3819 | 0.1195 | 61.7566 | 516.6535 | 25.5637 | 113.6351 |
| CC065/CC032 | 1 | 6.9248 | 0.6535 | 10.5966 | 16.2153 | 4.2346 | 19.8256 |
| CC001/CC005 | 2 | 3.5568 | 0.0856 | 41.5597 | 485.6076 | 10.1877 | 94.5724 |
| CC001/CC074 | 2 | 3.6335 | 0.1091 | 33.2942 | 305.0778 | 8.3261 | 75.2418 |
| CC002/CC074 | 2 | 3.1555 | 0.1246 | 25.3247 | 203.2434 | 5.5120 | 59.9535 |
| CC011/CC004 | 2 | 2.8864 | 0.0082 | 352.6076 | 43075.2195 | 69.6692 | 860.4619 |
| CC011/CC050 | 2 | 2.5690 | 0.0606 | 42.4150 | 700.2869 | 7.2981 | 107.8075 |
| CC012/CC004 | 2 | 2.8068 | 0.0502 | 55.8607 | 1111.7501 | 10.6902 | 137.6402 |
| CC012/CC016 | 2 | 2.3582 | 0.0096 | 245.3774 | 25532.1567 | 37.7787 | 643.1432 |
| CC012/CC041 | 2 | 3.8042 | 0.2583 | 14.7286 | 57.0247 | 3.8405 | 32.7989 |
| CC015/CC005 | 2 | 2.1236 | 0.0527 | 40.2684 | 763.5644 | 5.3449 | 109.6870 |
| CC015/CC011 | 2 | 1.4407 | 0.0113 | 127.3592 | 11258.3650 | 8.4105 | 403.3653 |
| CC021/CC002 | 2 | 1.8923 | 0.0232 | 81.6319 | 3521.5712 | 9.0411 | 232.2440 |
| CC021/CC032 | 2 | 3.0526 | 0.0443 | 68.8381 | 1552.3221 | 14.4659 | 164.8006 |
| CC023/CC005 | 2 | 1.9148 | 0.0121 | 158.3470 | 13094.8564 | 17.8825 | 448.4713 |
| CC023/CC042 | 2 | 3.9013 | 0.0821 | 47.5361 | 579.2110 | 12.6746 | 105.0137 |
| CC025/CC042 | 2 | 2.1192 | 0.0325 | 65.2451 | 2008.7430 | 8.6332 | 177.8599 |
| CC026/CC042 | 2 | 1.3012 | 0.0126 | 103.0813 | 8166.1341 | 5.3966 | 340.3386 |
| CC028/CC025 | 2 | 3.2856 | 0.1864 | 17.6224 | 94.5177 | 3.9980 | 41.1598 |
| CC030/CC028 | 2 | 2.4640 | 0.0494 | 49.8899 | 1010.1626 | 8.1401 | 128.7100 |
| CC065/CC003 | 2 | 2.4580 | 0.0245 | 100.4102 | 4101.8562 | 16.3313 | 259.2731 |
| SP 1 sum | 1 | 68.6338 | 2.8619 | 23.9819 | 8.3797 | 18.6461 | 29.9789 |
| SP 2 sum | 2 | 32.7237 | 1.2374 | 26.4462 | 21.3729 | 18.1730 | 36.2469 |
| Overall sum |  | 100.3575 | 4.0993 | 24.4818 | 5.9722 | 19.9271 | 29.4983 |

**Table S3.**
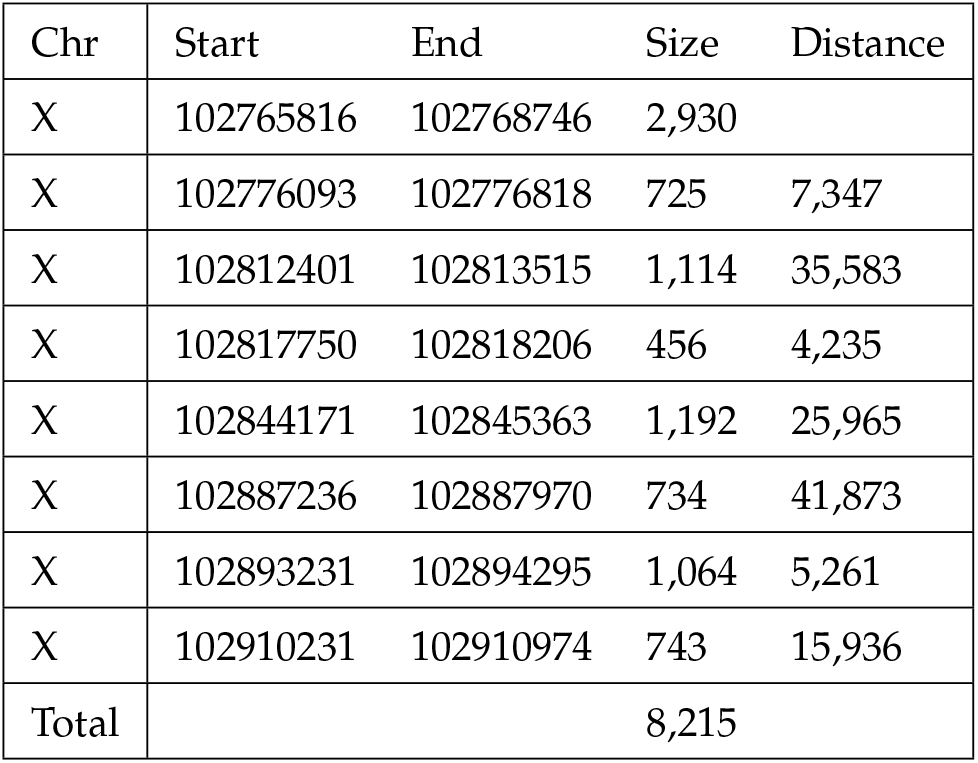
Assembled 8 sequences spanning the *Xce* interval generated from high-coverage CC WGS used to infer the phylogenetic relationships between the CC strains based on this region.

## Notes

### Competing Interest Statement

The authors have declared no competing interest.

