## Supplementary material for "Bayesian modeling of skewed X inactivation in genetically diverse mice identifies a novel *Xce* allele associated with copy number changes": FileS1.docx

**Skewed X-inactivation in genetically diverse mice is associated with recurrent copy number changes at the mouse *Xce* locus**

Kathie Sun, Daniel Oreper, Sarah Schoenrock, Rachel McMullan, Paola Giusti-Rodriguez, Vasyl Zhabotynsky, Darla Miller, Lisa Tarantino, Fernando Pardo-Manuel de Villena, William Valdar

File S1 (this file): Data documentation.

(1) R scripts for running all analysis and (2) additional intermediate data files (inferred haplotype blocks for SP1/2, summary-level results from Bayesian regression models, extended demographic information for samples in SP1/2, etc.) available at <https://github.com/kathiesun/XCI_analysis>

File S2: Demographic information on 266 mouse samples, and summary of results (CSV)

File S3: Data on 25-mers used to quantify gene expression (CSV)

Columns: Gene|Chromosome|rsId|Position|ProbeSeq|TranscriptID|Ref|Alt|A|B|C|D|E|F|G|H

A-H refer to the expected reference (0) or alternate (1) allele in each of the 8 CC founder strains

File S4: Posterior mode, mean, median, and 95% highest posterior densities determined by the

Bayesian hierarchical model for all covariates in GLM performed for SP1 and SP2 (CSV).

File S5: Counts of 45-mers from CC and CC-founder strains used for determining CNVs (CSV)

(columns: position | sequence | copy number in reference sequence | count of k-mer in each founder sample)

File S6: Counts of 25-mers centered at a reference allele from SP1 RNA-seq data (CSV)

(columns: sample | kmer | direction | count)

File S7: Counts of 25-mers centered at an alternate allele from SP1 RNA-seq data (CSV)

(columns: sample | kmer | direction | count)

File S8: All counts of 25-mers from SP2 RNA-seq data (zipped directory containing CSV files)

(columns: sample | kmer | direction | count)

File S9: Plots of XCI proportion estimates for all 266 samples (PDF)

File S10: Positions of seven CNVs (segmental duplications and inversions) in *Xce* (TXT)

Table S1: Summary of RIX crosses

Table S2: alpha0 posterior distributions

Table S3: Sequences used to infer phylogeny in CC strains

Figure S1: DAG in detail

Figure S2: Bionano optical mapping alignment

Figure S3: Counts of 45-mers spanning *Xce* in C57BL/6J

Figure S4: Counts of 45-mers spanning *Xce* in A/J

Figure S5: Counts of 45-mers spanning *Xce* in 129S1/SvlmJ

Figure S6: Counts of 45-mers spanning *Xce* in CAST/EiJ

Figure S7: Counts of 45-mers spanning *Xce* in PWK/PhJ

Figure S8: Counts of 45-mers spanning *Xce* in WSB/EiJ

Figure S9: Counts of k-mers spanning ends of SD7 for C57BL/6J and NZO
