## Supplementary material for "Bayesian modeling of skewed X inactivation in genetically diverse mice identifies a novel *Xce* allele associated with copy number changes": FigS9_NZObounds.pdf

a.

### CC Genome: CC002M4575_UNC_NYGC

**1,165,869,714 strings with 177,212,196,528 bases and index size of 22,844,202,515 bytes (1.03 bits per base)**

**Target: CAACTGAAGCTTCTTTCTCTGTGATAACTCCAGCTGTGTCAAGTTGACACAAAACTAGCCAGTACAGTAGGAA**

Found 29 times (12 forward, 17 reverse-complemented)

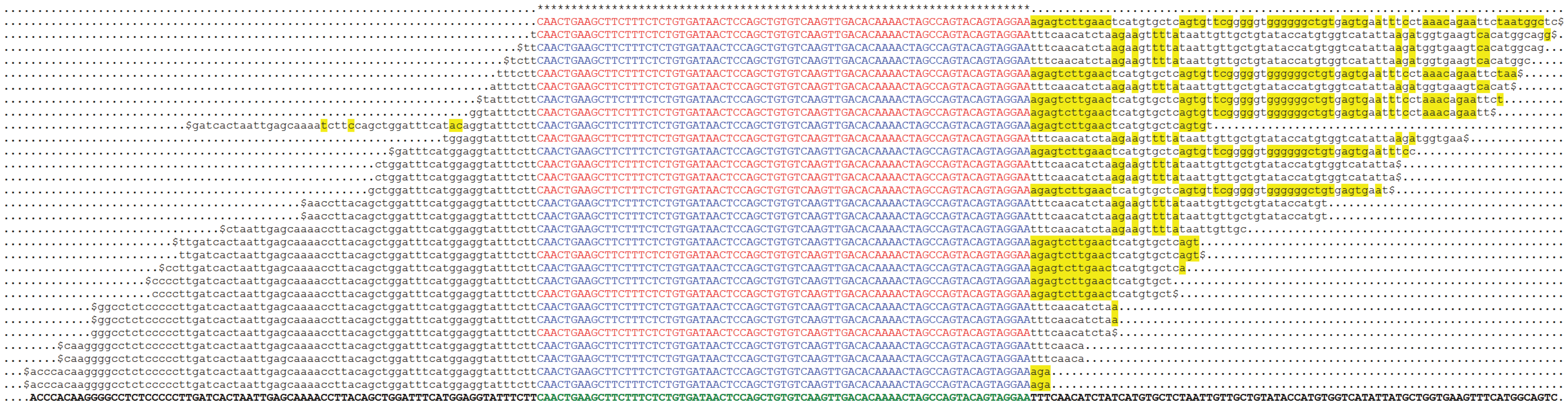

### CC Genome: CC005M4714_UNC_NYGC

**1,103,912,912 strings with 167,794,762,624 bases and index size of 21,818,642,651 bytes (1.04 bits per base)**

**Target: CAACTGAAGCTTCTTTCTCTGTGATAACTCCAGCTGTGTCAAGTTGACACAAAACTAGCCAGTACAGTAGGAA**

Found 13 times (6 forward, 7 reverse-complemented)

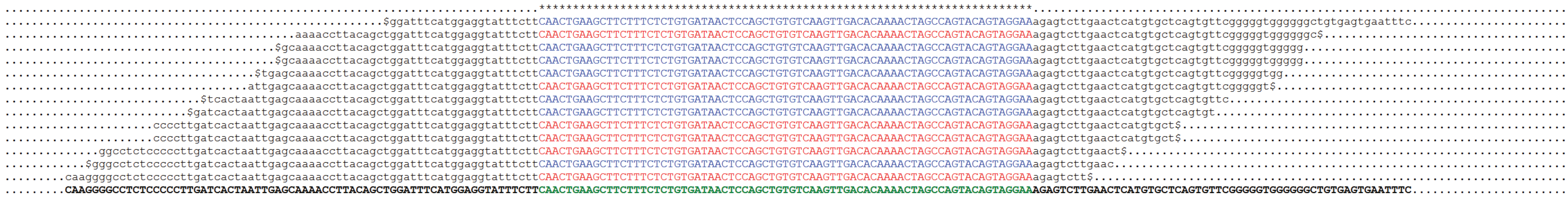

Results table ⊟

Show All entries | Show/hide columns (2 hidden) | X: S

| Genomic Location | Overlapping Gene(s) | Orientation | Query start | Query end | Length | Score | E-val | %ID |
|---|---|---|---|---|---|---|---|---|
| X:102842904-102843102 [Sequence] | | Forward | 1 | 199 | 199 [Sequence] | 388.0 | 3.6e-108 | 100.00 [Alignment] |

b.

### CC Genome: CC002M4575_UNC_NYGC

**1,165,869,714 strings with 177,212,196,528 bases and index size of 22,844,202,515 bytes (1.03 bits per base)**

**Target: GACTCATCCCCAACAGTAACTTGTCTCTATACTCTGGGGTCTATGAGGGCATCTGGCCAAT**

Found 25 times (18 forward, 7 reverse-complemented)

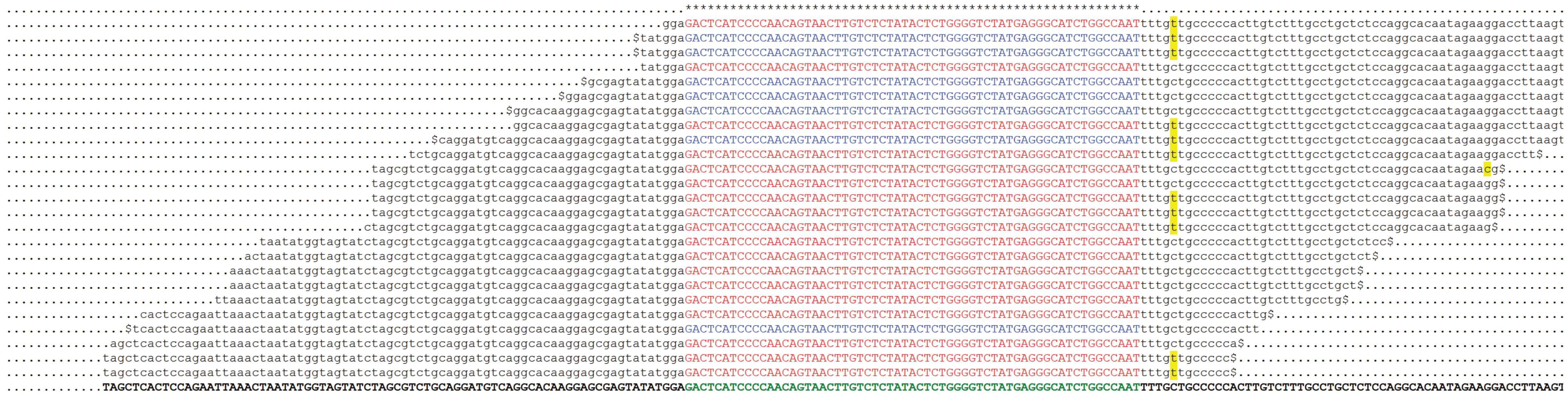

### CC Genome: CC005M4714_UNC_NYGC

**1,103,912,912 strings with 167,794,762,624 bases and index size of 21,818,642,651 bytes (1.04 bits per base)**

**Target: GACTCATCCCCAACAGTAACTTGTCTCTATACTCTGGGGTCTATGAGGGCATCTGGCCAAT**

Found 15 times (10 forward, 5 reverse-complemented)

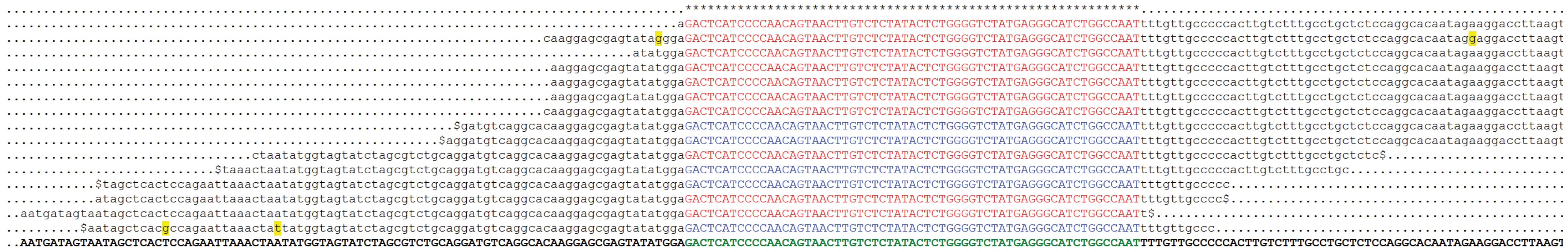

| Genomic Location | Overlapping Gene(s) | Orientation | Query start | Query end | Length | Score | E-val | %ID |
|---|---|---|---|---|---|---|---|---|
| X:102839320-102839380 [Sequence] | | Forward | 1 | 61 | 61 [Sequence] | 119.0 | 3.6e-27 | 100.00 [Alignment] |
| X:102802350-102802410 [Sequence] | | Forward | 1 | 61 | 61 [Sequence] | 117.0 | 2.0e-26 | 98.36 [Alignment] |
| X:102855166-102855226 [Sequence] | | Reverse | 1 | 61 | 61 [Sequence] | 117.0 | 2.0e-26 | 98.36 [Alignment] |
