## Supplementary figures and images for "Bayesian modeling of skewed X inactivation in genetically diverse mice identifies a novel *Xce* allele associated with copy number changes"

### FigS1_dag.pdf

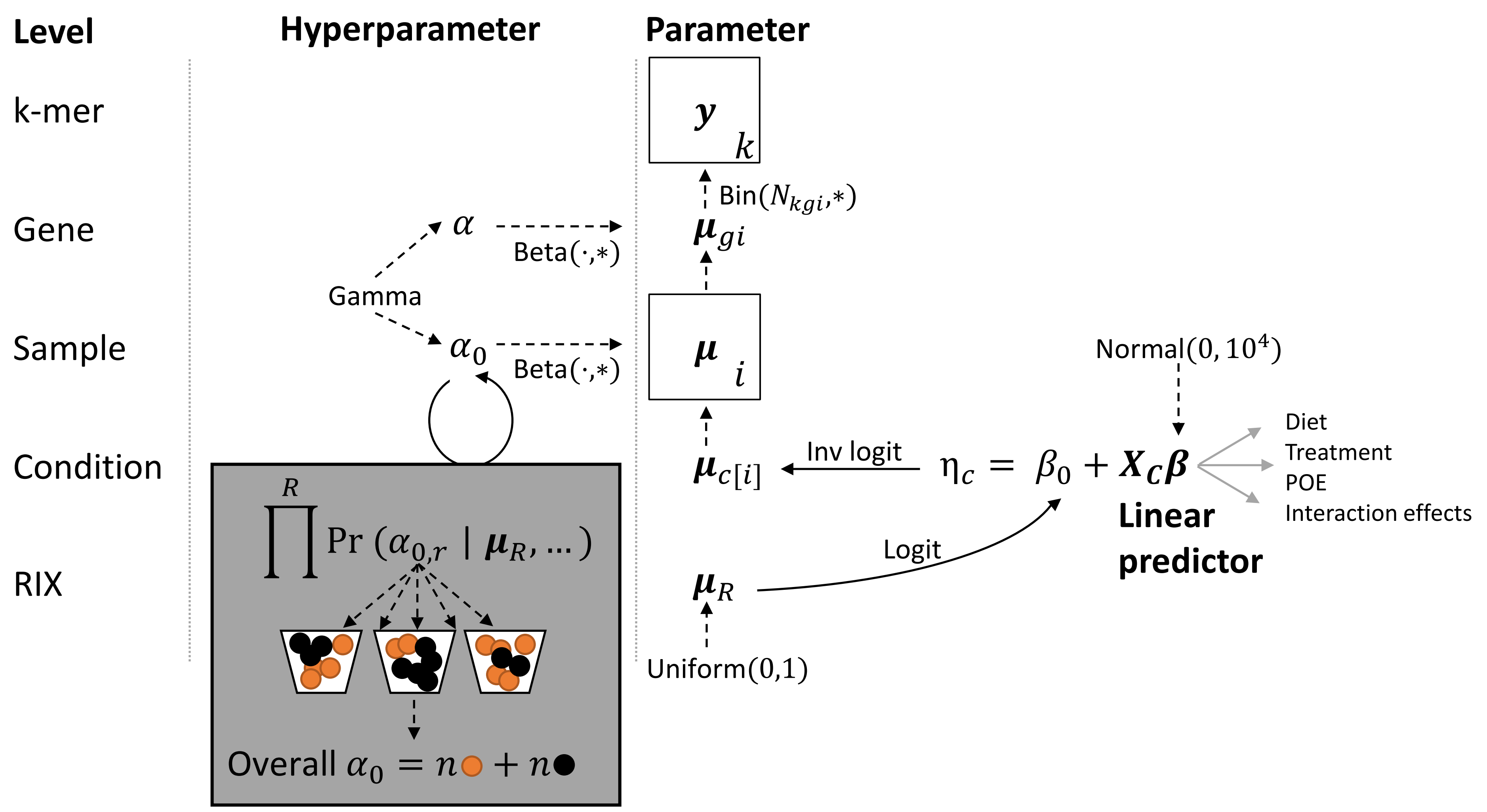

### FigS2_bionano_B6.pdf

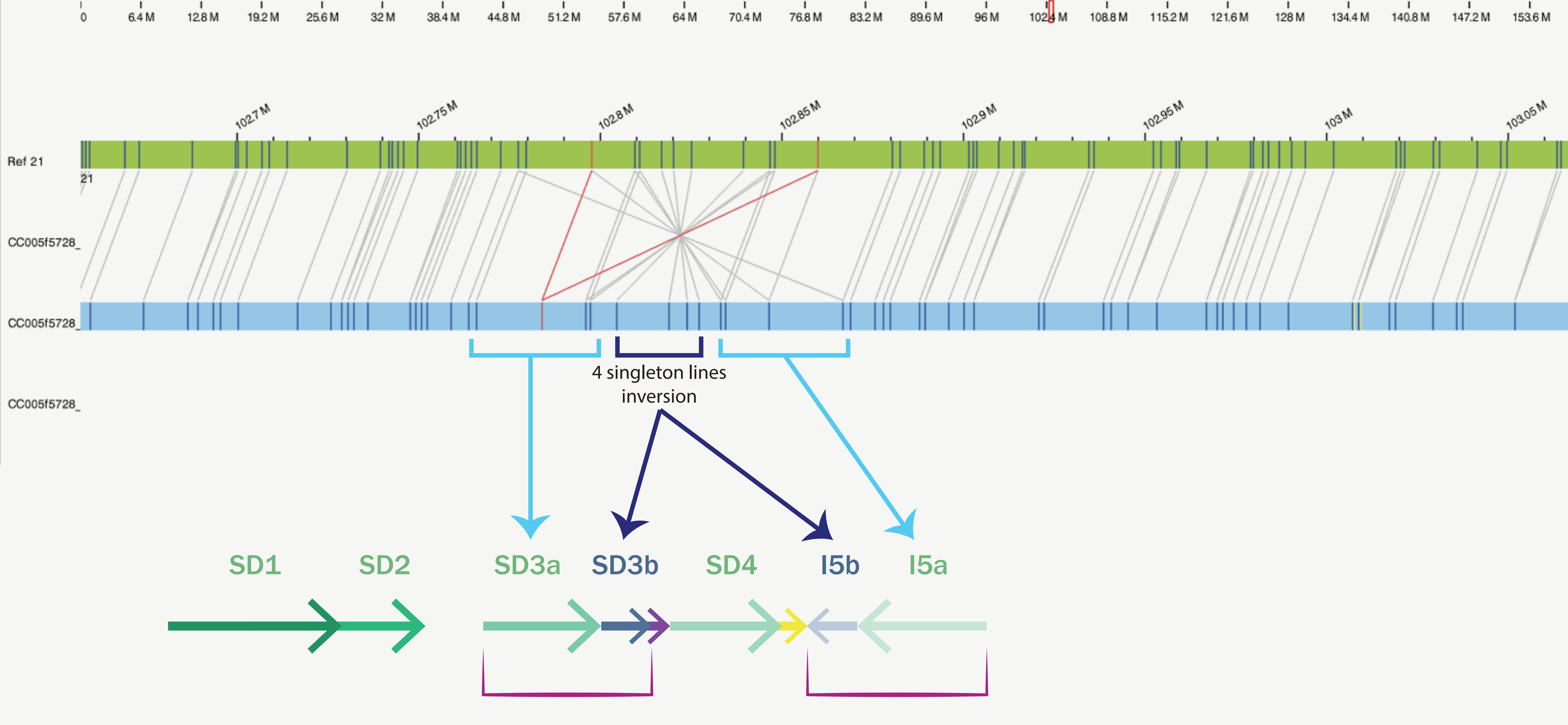

### FigS3_B6.pdf

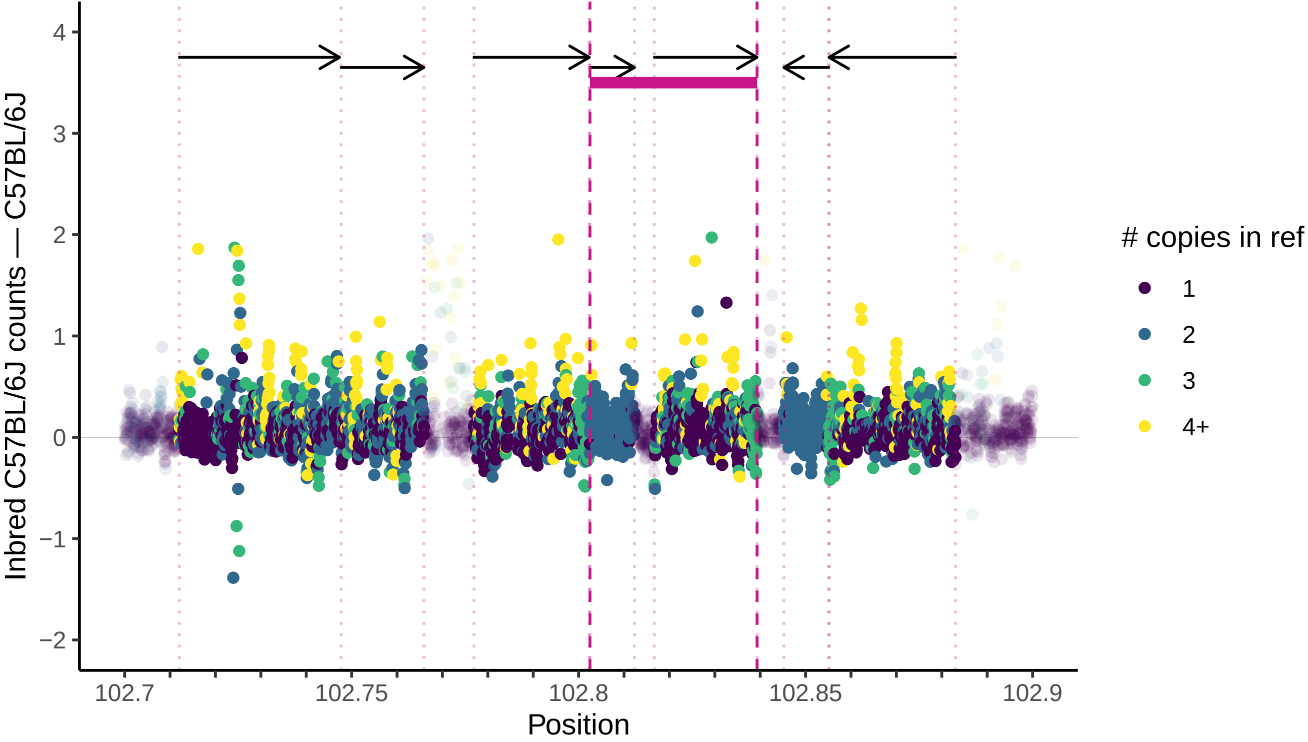

### FigS4_aj.pdf

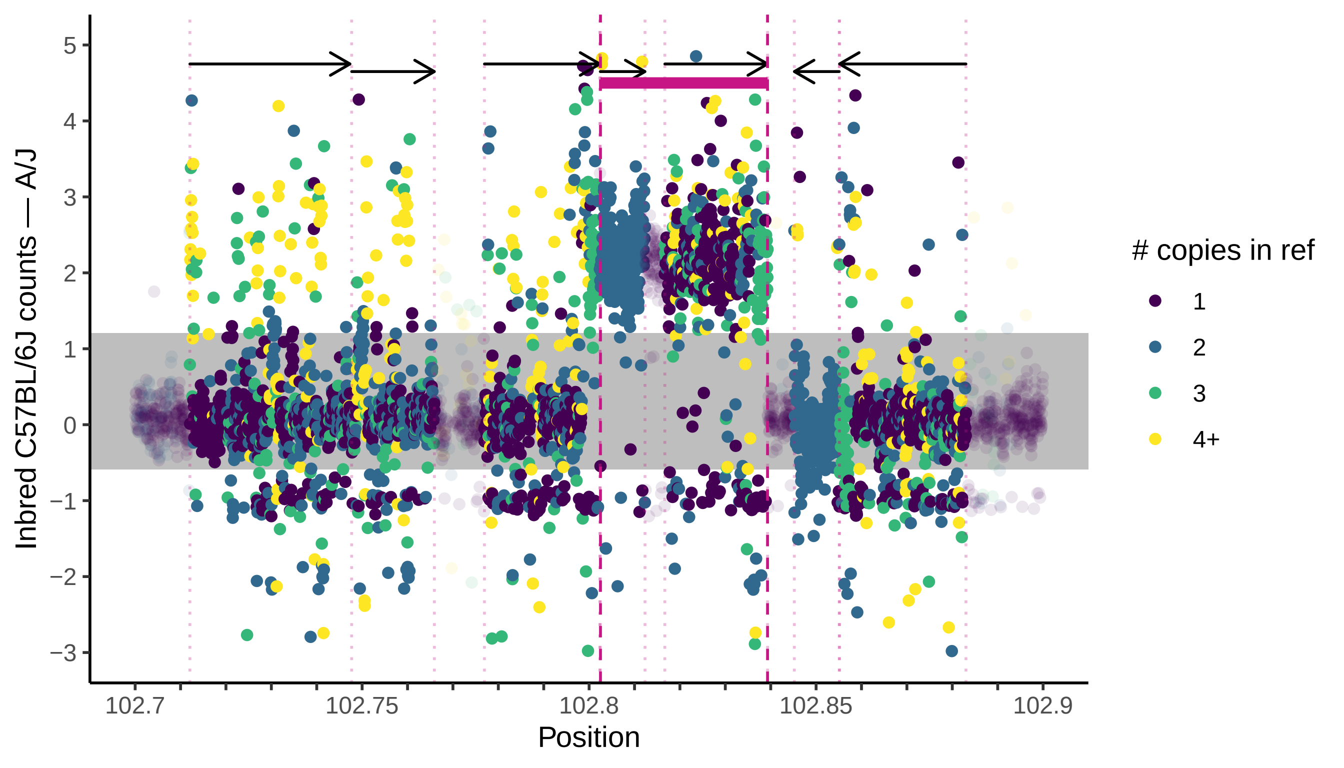

### FigS5_129.pdf

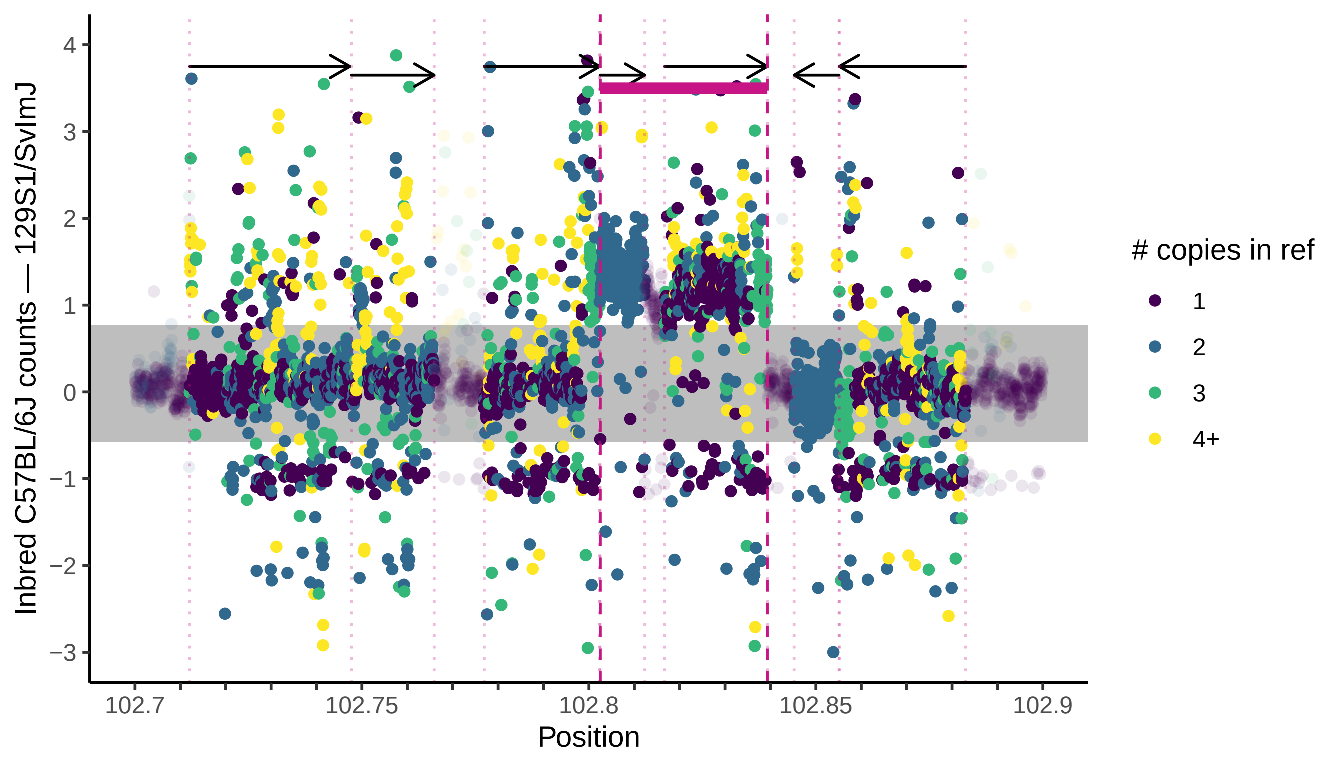

### FigS6_CAST.pdf

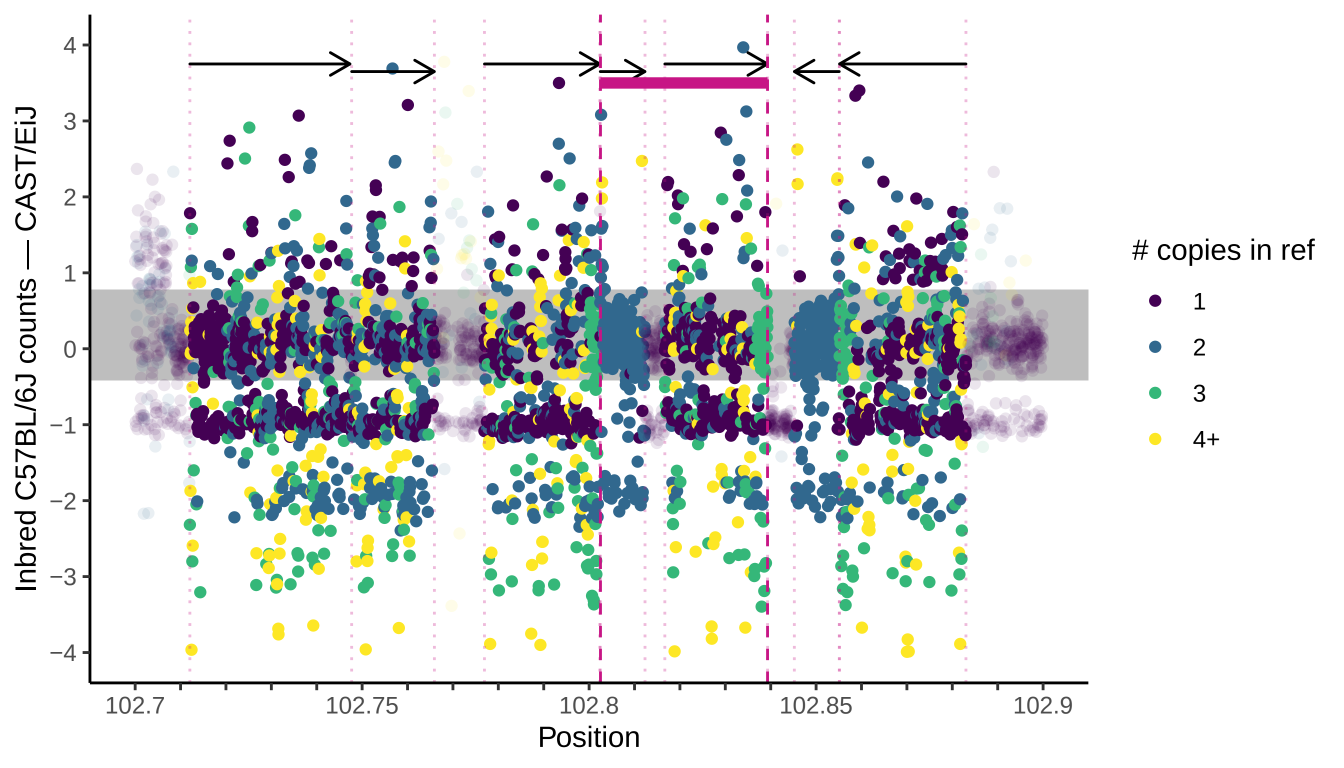

### FigS7_PWK.pdf

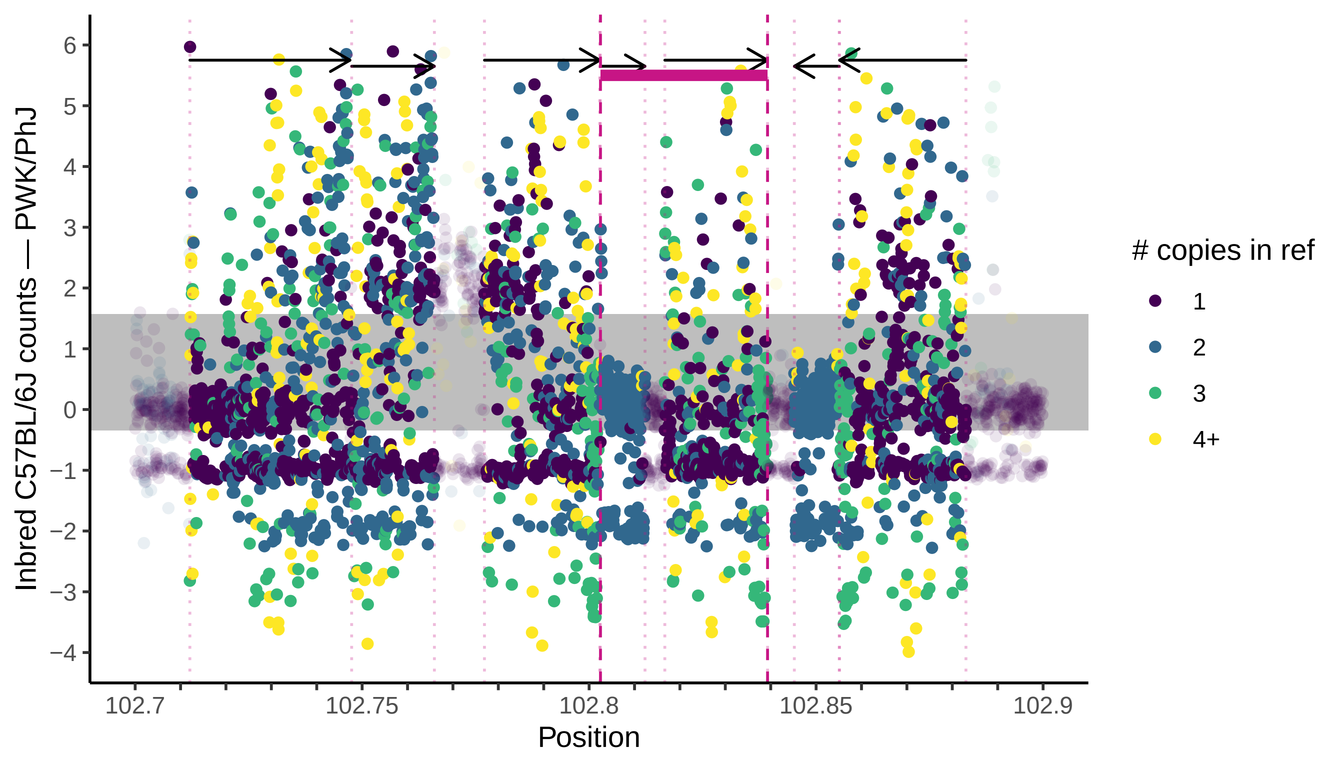

### FigS8_WSB.pdf

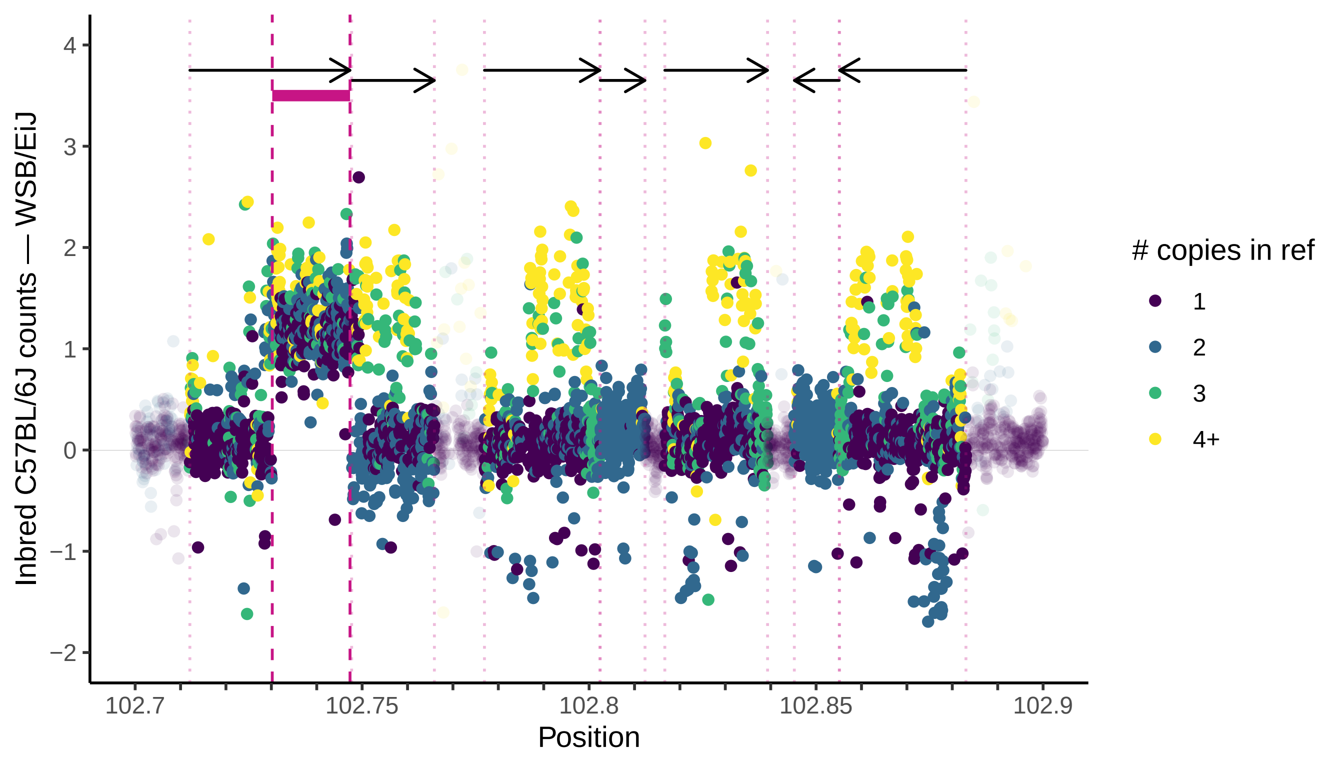
